# Long-lasting, subtype-specific regulation of somatostatin interneurons during sensory learning

**DOI:** 10.1101/2024.11.19.624383

**Authors:** Mo Zhu, Matthew B. Mosso, Xiaoyang Ma, Eunsol Park, Alison L. Barth

## Abstract

Somatostatin (SST)-expressing inhibitory neurons are a major class of neocortical γ-amino butyric acid (GABA) neurons, where morphological, electrophysiological, and transcriptomic analyses indicate more than a dozen different subtypes. However, whether this diversity is related to specific roles in cortical computations and plasticity remains unclear. Here we identify learning-dependent, subtype-specific plasticity in layer 2/3 SST neurons of the mouse somatosensory cortex. Martinotti-type, SST neurons expressing calbindin-2 show a selective decrease in excitatory synaptic input and stimulus-evoked calcium responses as mice learn a stimulus-reward association. Using these insights, we develop a label-free classifier using basal activity from *in vivo* imaging that accurately predicts learning-associated response plasticity. Our data indicate that molecularly-defined SST neuron subtypes play specific and highly-regulated roles in sensory information processing and learning.

---

Single-cell transcriptomic analyses have redefined the complexity of neocortical networks, where more than a hundred molecularly-distinct cell-types have been identified (*1*). The functional relevance of these cell types in cortical computation and plasticity is largely unexplored. In particular, somatostatin-expressing (SST) γ-amino butyric acid (GABAergic) neurons in the neocortex are highly heterogeneous, where studies have proposed an increasing number of distinct cell-types (from 6 in 2016 to nearly 40 in 2023; (*2*)). Because SST neurons are one of three major classes of cortical interneurons, densely wired into the cortical network, with robust responses that are modulated by both sensory input and cognitive variables, the functional diversity of this class is of particular interest.

In sensory neocortex, SST neurons exhibit broad receptive fields (*3*) and may encode cognitive information such as mismatch detection and attention (*4*, *5*). Dynamic modulation of layer 2/3 (L2/3) SST Ca^++^ responses during task performance has been reported in both sensory and motor cortex (*6–8*), and SST neuron activity has been implicated in some forms of learning (*9*, *10*). Heterogeneous SST responses, even in superficial layers, have confounded efforts to detect significant changes in population activity across conditions (*8*, *11–13*). Thus, mechanisms driving potential response plasticity and whether this plasticity might be differentially expressed across transcriptionally-distinct subtypes of SST neurons is unclear. Without this information, it is difficult to advance hypotheses about the role of SST neurons in cortical computations and learning.

## Stimulus-reward coupling drives SST response plasticity *in vivo*

We initially sought to investigate long-lasting changes in SST neurons that were associated with learning, using a home-cage, freely-moving training paradigm where the contingency of sensory stimuli and reward outcomes could be easily manipulated in a whisker-dependent learning task (*14*). For longitudinal *in vivo* imaging of neural activity, GCaMP6f was expressed in SST neurons in SST-Cre x Ai148 transgenic mice and a cranial window was implanted over primary somatosensory (barrel) cortex (Fig. 1A). Animals were acclimated to the homecage training environment for 6 days before training onset, where they could freely initiate trials for water that was delivered at a recessed lickport without any predictive cue (80% probability). During sensory association training (SAT), a gentle multiwhisker stimulus (4-6 psi airpuff) preceded water delivery for 80% of trials, and 20% of trials had no stimulus and no reward (blank trials; Fig. 1A and fig. S1). Learning was assessed by comparing the frequency of anticipatory licking (300 ms prior to water delivery) to stimulus trials and blank trials, and the majority of animals showed an increase in anticipatory licking to stimulus trials after 1-2 days of training (fig. S1). Stimulus-evoked Ca^++^ transients in SST neurons were imaged in head-fixed animals outside of the training context, to avoid confounds introduced by motivational state, licking and other goal-directed movements, and cognitive variables related to performance accuracy.

**Figure 1:**
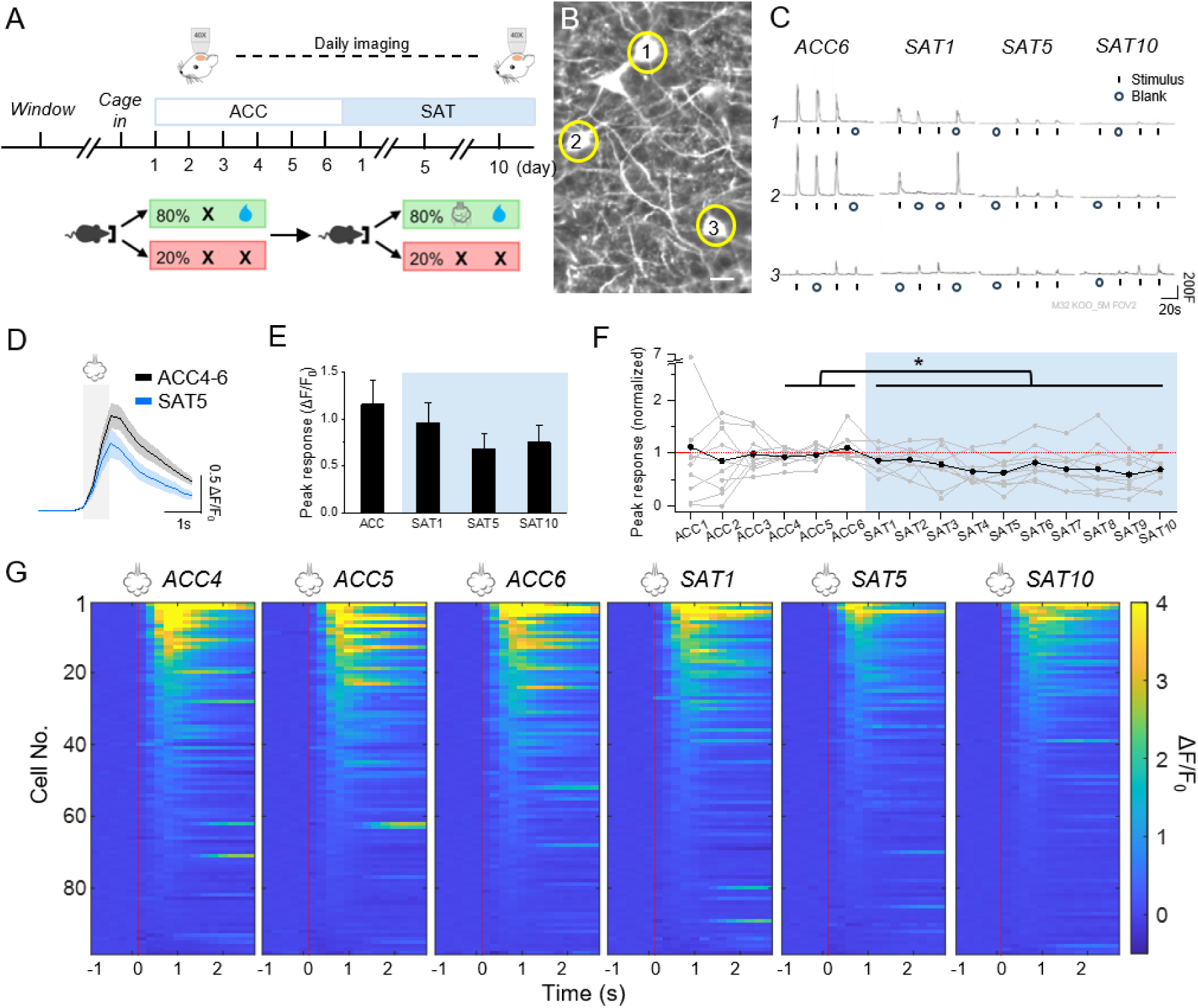
Sensory association training (SAT) reduces stimulus-evoked responses in SST neurons. (**A**) Top: experimental timeline for longitudinal calcium imaging. Bottom: Structure of sensory association training. (**B**) An example imaging FOV. Scale=20 µm. (**C**) Example traces of airpuff-evoked responses from the same example cells on ACC6, SAT1, SAT5, and SAT10. F (fluorescence) is reported in arbitrary units. (**D**) Mean ± SEM airpuff-evoked response during acclimation 4-6 (ACC4-6) and SAT5. Animal average, N=10 mice. Grey shade indicate the airpuff period. (**E**) Mean of evoked response peak on ACC4-6, SAT1, SAT5, and SAT10 by mice. Paired t-test with Bonferroni correction, comparing ACC4-6 with training days, p=0.18, 0.061, and 0.25 for SAT1, SAT5, and SAT10, respectively. (**F**) Peak airpuff-evoked response across the acclimation and training period, averaged across mice and normalized to ACC4-6 (black; n=98 cells in 10 mice). Mean ± SEM of shown in the figure. Grey lines indicate mean cell response for individual mice. One-way repeated measures ANOVA, p=2.3×10^−5^. (**G**) Heatmaps of the evoked response traces of all cells rank ordered within individual imaging days. Red dotted lines indicate airpuff onset.

Daily imaging of L2/3 SST neurons in barrel cortex across 6 days of pretraining acclimation (ACC) to the training environment and 10 days of SAT revealed a progressive reduction in mean sensory-evoked Ca^++^ activity that was initiated at the onset of training (Fig. 1B-G; animal average ΔF/F_0_, ACC 1.2±0.3 vs. 1 day training (SAT1) 0.96±0.2, SAT5 0.68±0.2, and SAT10 0.75±0.2; n=98 cells in 10 mice; ACC4-6 vs SAT1-10 p=2×10^−5^ by ANOVA; see fig. S2 for histological verification of imaging sites). This response suppression was not observed in barrel cortex from animals subjected to pseudotraining (PSE; fig. S3) where stimulus probability (80% of trials) was preserved but uncoupled to reward; indeed SST responses were modestly elevated, an increase that was not significant (fig. S4: animal average ΔF/F_0,_ ACC 0.82±0.3 vs. PSE1 1.0±0.1, PSE5 1.4±0.3, PSE10 1.6±0.6; n=62 cells in 5 mice; ACC4-6 vs PSE1-10 p=0.074 by ANOVA; see fig. S5 for histological verification). No training-induced changes in SST neurons were observed in higher-order cortical areas (fig. S6). No sex differences were observed for either group (fig. S7).

Long-lasting changes in SST activity in somatosensory cortex suggest that information-encoding circuits during learning might be saturated by our training paradigm. To investigate this, after 10 days of SAT animals were retired to their home cage where training was discontinued. Importantly, SST population activity returned to baseline levels, at least after several weeks of housing in conventional animal caging (ΔF/F_0_ ACC6 0.36+0.12, SAT10 0.19+0.06, post-training homecage 0.52+0.14, n=29 cells in 3 mice; p=0.12 by ANOVA). Thus, SST activity suppression initiated by sensory learning may require sustained training in order to be maintained.

Because the same SST neurons could be identified across days, response profiles for individual neurons could be aligned across the imaging period (Fig. 1). Daily tracking of individual SST neurons revealed that responses to sensory stimulation were highly heterogeneous (100-fold variation between neurons). Furthermore, response plasticity during SAT was also diverse, with some SST neurons decreasing and others increasing stimulus-evoked Ca^++^ transients. These properties were characteristic of individual neurons and were also maintained across the training period (fig. S8). Although heterogeneous response properties of SST neurons in superficial layers of sensory and motor cortex have been previously described (*7*, *8*, *15*, *16*), it has not been easy to connect these dynamic features to the molecular phenotype of putative SST subtypes (but see (*17*)). The expansion of SST neuronal subtypes defined by gene expression (*1*) has suggested functional differentiation, but evidence for this has been limited (*18*).

## Anatomical synaptic analysis reveals subtype-specific SST plasticity

We hypothesized that the suppression of SST sensory-evoked responses, particularly as observed outside of the training context, might be linked to long-lasting changes in SST neurons during SAT (14, 19). Intrinsic excitability of SST neurons was similar before and after training (fig. S9). Because they are densely connected into the local network of pyramidal neurons (*20*), we hypothesized their decreased activity might be related to a reduction in excitatory synaptic drive. For quantitative analysis of excitatory synaptic input in fixed tissue, a Cre-dependent PSD95-FingR intrabody (*19*) tagged with mCitrine was virally transduced in barrel cortex for SST synaptic labeling (Fig. 2A-E). Synaptic PSD95 levels are directly correlated with excitatory synaptic strength, and are bidirectionally modified during synaptic potentiation or depression (*21*, *22*). Thus, tracking synaptic PSD95 can yield insight into bidirectional excitatory plasticity. Automated detection and volume analysis of thousands of PSD95 puncta from L2/3 SST neurons (fig. S10) showed a small but significant population decrease in volume between ACC and SAT conditions, particularly at later stages of training (fig. S11A-C).

**Figure 2.**
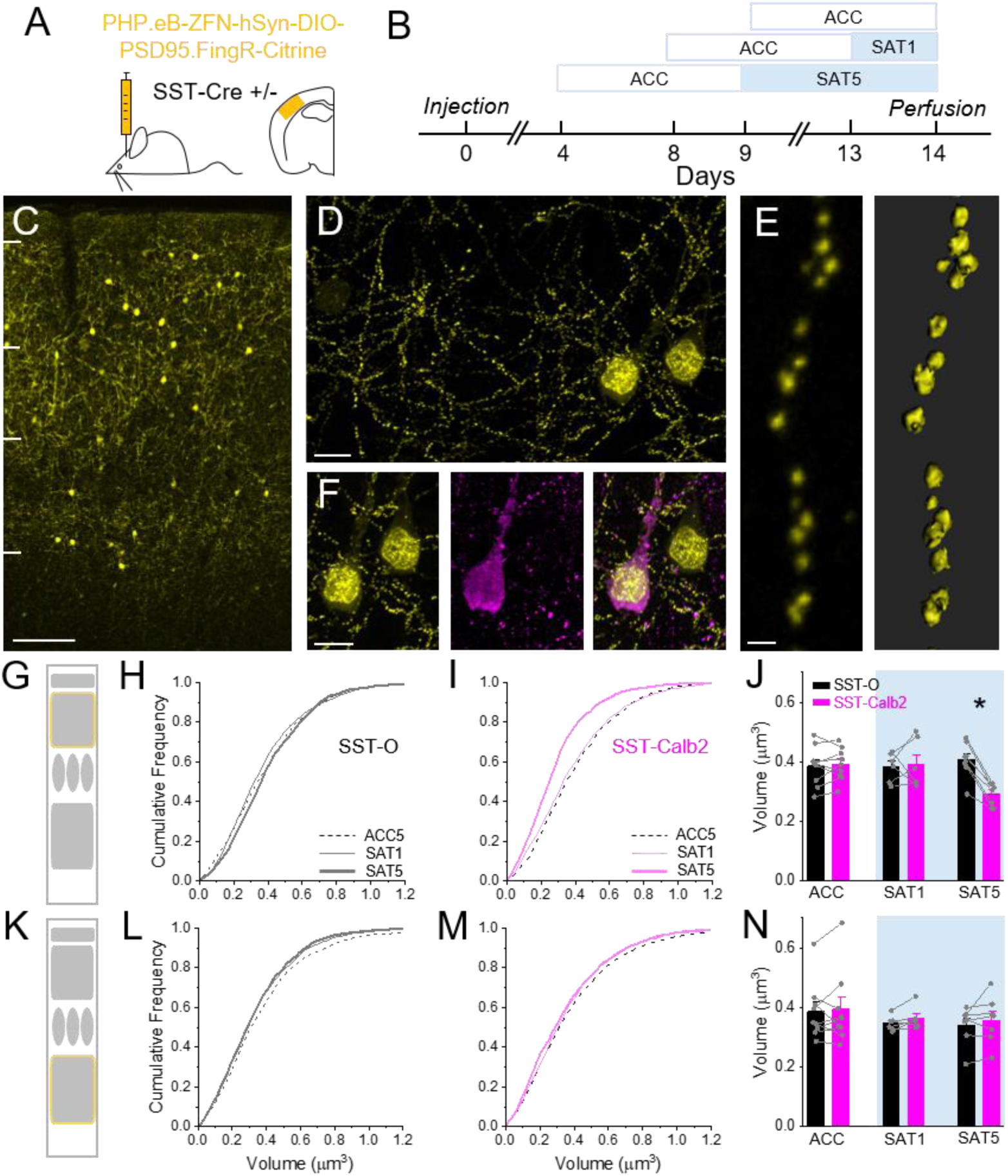
Excitatory synapses in SST-Calb2 neurons are selectively suppressed after 5 days of sensory association training. (**A**) DiO-PSD95.FingR-Citrine was injected in barrel cortex of SST-Cre mice. (**B**) Schematic outlining injection and training timeline. Animals in fixed tissue analysis were time matched to control for viral expression time. (**C**) 10x image of SST-Cre mice expressing PSD95.FingR across the cortical column. Scale=100µm (**D**) 63x volumetric image of PSD95.FingR labeled somatostatin neurons. Scale=10µm (**E**) (Left) Zoomed image of PSD95.FingR puncta along a dendrite. (Right) Surface reconstruction of citrine signal. Scale=1µm (**F**) Immunohistochemical labeling of SST neurons (Left) PSD95.FingR-Citrine (Middle) calretinin expression (protein product of Calb2 gene) Scale=10µm (Right) merged channels. (**G**) Schematic indicating layer imaging field of view was captured (Layer 2/3). (**H**) Cumulative distribution of PSD95 puncta volume in layer 2/3 SST-O neurons. (**I**) Same as (H) for L2/3 SST-Calb2 neurons. (**J**) Within animal comparison of mean PSD95 puncta volume between SST-O (gray bars) and SST-Calb2 (pink bars) after experiencing acclimation (ACC), one, or five days of sensory association training (SAT). (**K-N**) Same as G-J but for L5 SST-O and SST-Calb2 SST neurons.

L2/3 SST neurons in barrel cortex are anatomically and molecularly heterogeneous, comprised of SST-Calb2 (Martinotti type, with a prominent axon in L1), SST-Mme (targeting mainly L2/3), and SST-Chodl neurons (long-range projecting) (*23*, *24*). Recent transcriptomic analysis suggests that there may be at least several other subtypes contained within these three subclasses (*1*). We hypothesized that the modest shift in PSD95 puncta size in L2/3 might be reflected in a subclass-dependent regulation of excitatory input strength. To investigate this, we immunostained PSD95-FingR SST neurons with an antibody directed against calretinin, the protein coded by the Calb2 gene (Fig. 2 F). PSD95 puncta size was compared between SST-Calb2 and unlabeled SST neurons (SST-other/SST-O) dendrites from control (ACC) and also trained mice (Fig. 2 G-J and K-N). In ACC mice, a comparison of PSD95 puncta in SST-Calb2 and SST-O dendrites showed no difference in puncta volume distribution (Fig. 2 H-J; ACC SST-O 0.39±0.02 vs. SST-Calb2 0.39±0.02 μm^3^, n=1,800 puncta in 9 mice). However, 5 days of SAT drove a prominent and highly significant reduction in puncta volume that was restricted to L2/3 SST-Calb2 dendrites (Fig. 2 H-J; SST-O 0.41±0.02 vs. SST-Calb2 0.29±0.01 μm^3^, n=1,600 puncta in 8 mice; paired t-test p=8×10^−4^). This difference was observed when averaged across animals and also when comparing adjacent SST-Calb2 and SST-O cells within the same field-of-view, where imaging conditions were matched (Fig. 2 J).

Because analysis was carried out in fixed tissue, we could also examine SAT-associated changes in PSD95 puncta size for SST-Calb2 cells in deeper layers. However, we did not observe a reduction in PSD95 puncta volume in L5, either overall (fig. S11 G-I) or for SST-Calb2 neurons in particular (Fig. 2 K-N; SAT5 SST-O 0.34±0.02 vs. SST-Calb2 0.36±0.03 μm^3^). Notably, a reduction in PSD95 puncta volume, either for SST-Calb2 or SST-O neurons, was not evoked by pseudotraining where whisker stimulation had no value in predicting water delivery (fig. S12 A-H).

Finally, to test whether active sensation during exploratory behaviors could be sufficient to induce changes in PSD95 puncta volume in SST-Calb2 neurons, mice were housed in an enriched environment for multiple days (fig. S12 I,J). However, this treatment also failed to modify either overall PSD95 puncta volume between SST-O and SST-Calb2 neurons, or the relative difference between SST-Calb2 and SST-O neurons within barrel cortex tissue from the same animal. Thus, the pairing of fully-predictive whisker stimulation and water reward during SAT uniquely drove a long-lasting reduction in excitatory input onto SST-Calb2 neurons. The relative timing of depressed stimulus-evoked SST activity (early during training) and a reduction in puncta volume (later in training) suggested that decreased SST neuron activity might initiate their postsynaptic reduction in PSD95 puncta volume. To test this hypothesis, we used chemogenetic suppression of activity by virally transducing the inhibitory DREADD receptor hM4Di into SST neurons in barrel cortex, in the absence of any training paradigm (fig. S13). Five days of CNO administration was sufficient to drive a reduction in PSD95 puncta in hM4Di-expressing SST neurons compared to unlabeled neurons in the same L2/3 field of view, for both SST-O and SST-Calb2 cells (fig. S13 C-H: SST-O comparison for hM4Di+ 0.29 ±0.01 vs. non-expressing SST-O 0.38±0.03 μm^3^, n=1,200 puncta in 6 mice each; paired t-test, p=0.01; fig. S13 I-N: SST-Calb2 comparison for hM4Di+ 0.23±0.01 vs. non-expressing SST-Calb2 0.36±0.02 μm^3^, n=1200 puncta in 6 mice each; paired t-test, p=0.004). Thus, chemogenetic suppression of activity in the broader population of L2/3 SST neurons could phenocopy the effects of SAT on PSD95 puncta size. The selective reduction in excitatory inputs onto SST-Calb2 neurons during SAT suggests these neurons are specifically engaged by neural circuits related to learning.

## SST-Calb2 neurons show selective response suppression during SAT

Are SST-Calb2 neurons responsible for the population decrease in sensory-evoked activity in L2/3 SST neurons observed during *in vivo* Ca^++^-imaging? *Calb2* is also expressed in a subpopulation of vasoactive intestinal peptide (VIP) interneurons, complicating efforts to monitor these neurons without intersectional genetics. Thus, we generated *SST-Flp* x *Calb2-Cre* mice with viral transduction of SST neurons using a Flp-dependent GCaMP6f and a Cre-dependent mCherry construct for detection of Calb2 neurons (Fig. 3A,B). Overlap in GCaMP6f and mCherry signal was an indicator of SST-Calb2 neurons (fig. S14), although it is important to note that *Calb2* is expressed either transiently or at low levels in other L2/3 SST subtypes (*23*, *24*), and post-hoc immunolabeling of Calb2 in SST neurons detects only ∼75% of genetically-labeled neurons from this intersectional cross (*25*), indicating that *Calb2* gene expression is not a precise indicator of protein levels. In general, Ca^++^ signals using virally-transduced GCaMP6f expression were weaker than observed in the transgenic Ai148 line, but both spontaneous and stimulus-evoked activity could be clearly distinguished.

**Figure 3.**
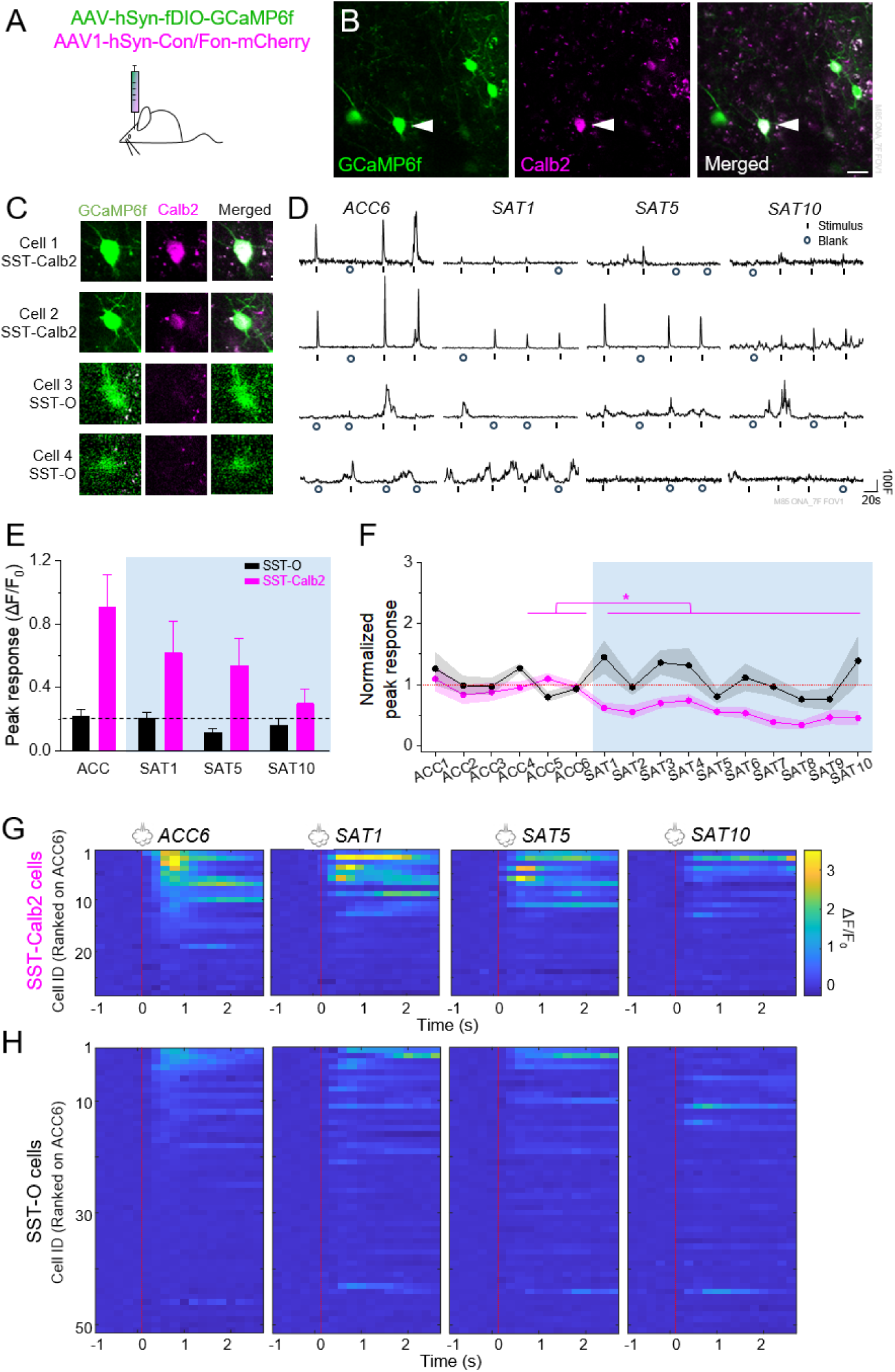
SAT selectively reduces whisker-evoked responses in SST-Calb2 neurons. (**A**) Schematic demonstrating the stereotaxic delivery of viral constructs into mouse barrel cortex. (B) Example FOV expressing GCaMP6f in SST cells and mCherry in Calb2-positive cells. Scale bar = 20 μm. (**C**) Left: example SST-Calb2 and SST-O neurons showing GCaMP6f expression (green) and Calb2 expression (magenta), with merged images (bottom) displaying both markers. (**D**) Example traces of example SST-Calb2 and SST-O cells on ACC6, SAT1, SAT5, and SAT10. (**E**) Mean peak response across ACC1-6 and SAT1-10. The magenta line represents the average peak response across 28 SST-Calb2 cells while the black line represents the average peak response across 51 SST-O cells collected in 7 mice. One-way repeated measures ANOVA, p=1.3×10^−12^ for SST-Calb2 and p=0.50 for SST-O. (**F**) Mean peak response of SST-Calb2 and SST-O cells on ACC4-6, SAT1, SAT5, and SAT10. Paired t-test with Bonferroni correction, comparing between SST-Calb2 and SS-O cells within different imaging days, p= 5.5×10^−5^, 0.10, 0.0018, and 0.12 on ACC4-6, SAT1, SAT5, and SAT10, respectively. (**G**) Response heatmaps of SST-Calb2 rank ordered on ACC6. Red line indicates in the onset of airpuff. (**H**) As in (G), but for SST-O cells.

Based on decreased PSD95 puncta volume in our fixed-tissue analysis, we predicted that SST-Calb2 neurons in barrel cortex would selectively show a reduction in stimulus-evoked activity across the SAT period (see fig. S15 for behavior and fig. S16 for histological verification). This was indeed the case (Fig. 3 C-H). SST-Calb2 neurons showed a significant reduction in peak amplitude of the whisker-evoked response that progressively decreased over training (SST-Calb2 fold-change: SAT1 0.62±0.083, SAT5 0.56±0.068, SAT10 0.46±0.12; n=28 cells in 7 mice; ACC4-6 vs. SAT1-10 p=1.3×10^−12^ by ANOVA). In comparison, unlabeled cells (which could represent SST-O neurons or SST-Calb2 neurons that were not virally transduced by the mCherry tag, designated for clarity as SST-O) showed no significant decrease in sensory-evoked activity (SST-O fold-change: SAT1 1.4±0.27; SAT5 0.81±0.12; SAT10 1.4±0.40; n=51 cells in 7 mice; ACC 4-6 vs SAT1-10 p=0.5 by ANOVA). These data indicate that sensory-evoked activity in SST-Calb2 neurons is selectively depressed by SAT, initiated prior to changes in excitatory input.

## Predictive classification of SST-Calb2 plasticity using *in vivo* Ca^++^ signals

Sensory-evoked activity in SST neurons is highly heterogeneous even under basal, homecage conditions without training (*12*, *15*). We observed that during ACC, SST-Calb2 neurons identified by mCherry expression typically showed a higher sensory-evoked response compared to unlabeled SST neurons (Fig. 4A). These data suggested that the SST-Calb2 subset of neurons might be defined by activity properties, irrespective of their plasticity during learning. Using 22 features of stimulus-evoked activity from this dataset (mainly related to response probability and peak stimulus-evoked responses over different time intervals; fig. S17 and S18), we applied clustering-informed heuristic classification to distinguish SST-Calb2 from SST-O neurons (Fig. 4B). Spontaneous activity was not used as a feature for classification, since it degraded cluster separation (fig. S19). Although the clustering did not exclusively assign SST-Calb2 neurons to one group, we reasoned that low levels of Calb2 expression in other SST subclasses (*1*) and also variability in viral uptake might account for this distribution. Notably, this classifier detected two other clusters, potentially reflecting the diversity of SST neurons in L2/3 (*24*, *25*).

**Figure 4.**
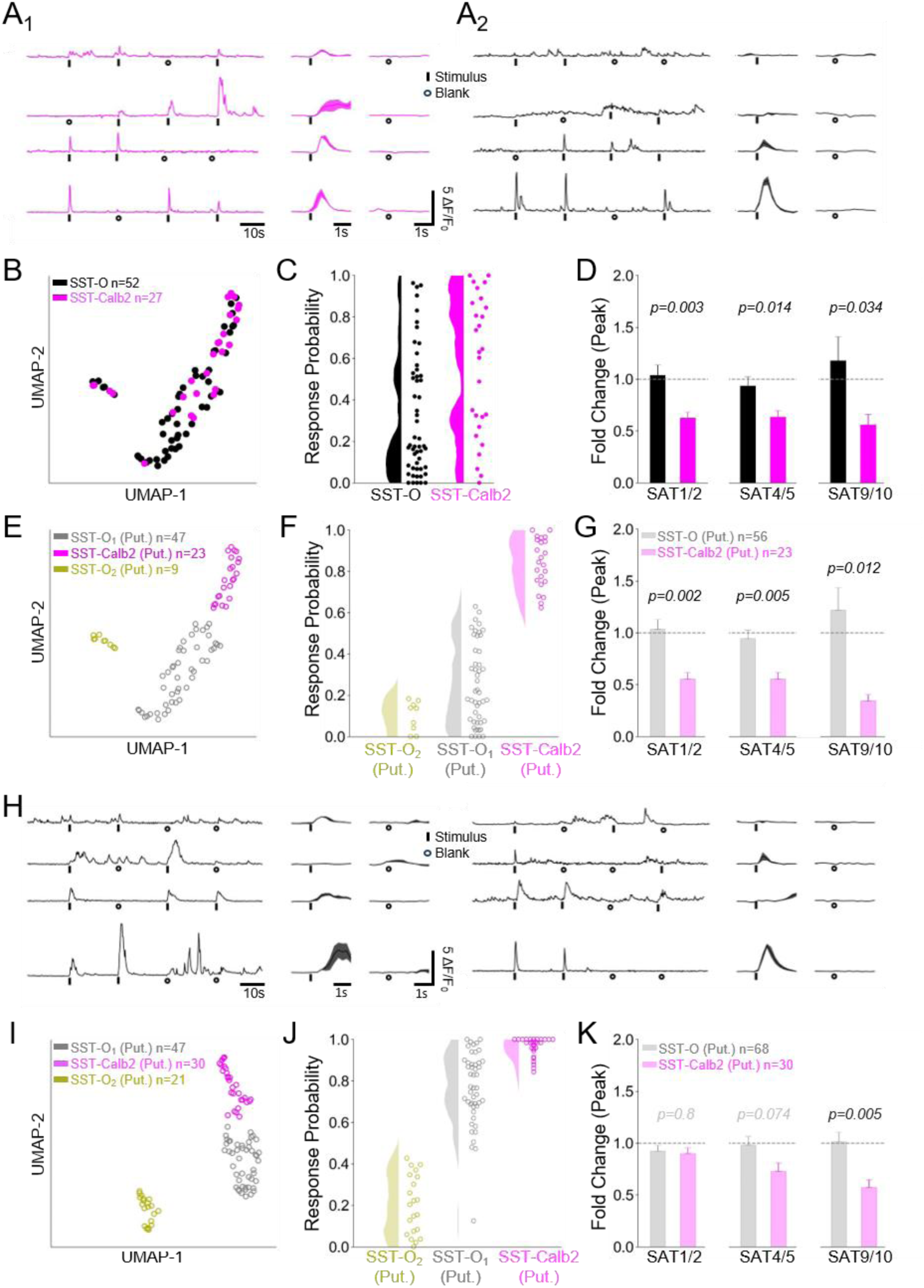
SST response features during ACC enable cluster-based classification that predicts response plasticity during learning. (**A**) Example Ca^++^ transients from SST-Calb2 neurons identified using SST-Flp x Calb2-Cre mice with viral labeling (see Fig. 3) during the pre-training stage (ACC5). (**A_2_**) As in A_1_ but for SST-O neurons. (**B**) UMAP visualization of SST neuron clustering using response properties for ACC4-6 (n=79 cells in 7 animals). (**C**) Example feature comparison of stimulus-evoked response probability for genetically-labeled SST-Calb2 and SST-O neurons. (**D**) SST-Calb2 neurons show significant reduction in mean peak evoked-response (all trials) during SAT, normalized to ACC4-6. Data averaged for SAT1/2, SAT5/6, and SAT9/10. (**E**) Spectral clustering assignment of putative SST-Calb2 (13/23 SST-Calb2 cells in the cluster) and putative SST-O neurons (10/47 and 3/9 SST-Calb2 cells) from the UMAP visualization in (B). (**F**) As in (C) but for the assigned, putative SST-Calb2 and combined putative SST-O groups. (**G**) As in (D), but for putative SST-Calb2 neurons versus SST-O_1_ and SST-O_2_ groups (combined for arison). Putative SST-Calb2 cells show significant reduction in mean peak evoked-response (all trials) during normalized to ACC4-6. (**H**) Example Ca^++^ transients from SST neurons during the pre-training stage (ACC5) the unlabeled, SST-Cre x Ai148 dataset (see Fig. 1). (**I**) As in (E) but for the unlabeled dataset, with putative ps identified by spectral clustering (n=98 cells). The putative SST-Calb2 group (30/98 cells) and putative SST-O 8 and 21/98 cells) were identified using the spectral clustering algorithm. (**J**) As in (F), but for clusters identified in nlabeled dataset. (**K**) As in (G), but for putative SST-Calb2 and combined putative SST-O groups from the eled dataset. All statistical comparisons done using a two-sample t-test.

Using this experimental grouping, we manually assigned all SST neurons in one cluster as putative SST-Calb2 neurons and combined the other clusters into a single group (putative SST-O; Fig. 4E). We then asked whether SAT-dependent response suppression would be concentrated in this SST-Calb2 subset. Remarkably, clustering based on SST activity properties prior to training (during ACC) showed strong predictive value in identifying neurons with SAT-dependent stimulus-evoked response suppression (Fig. 4G). Mean peak response from early (SAT1/2), mid (SAT4/5), and late (SAT9/10) timepoints was significantly reduced for putative SST-Calb2 classified neurons but not for putative SST-O classified neurons (putative SST-Calb2 fold-change vs. ACC4-6 for SAT1/2 0.554 ± 0.063; SAT5/6 0.552 ± 0.066; SAT9/10 0.343 ± 0.06; n=23 cells; putative SST-O fold-change versus ACC4-6 for SAT1/2 1.035 ± 0.093; SAT5/6 0.945 ± 0.082; SAT9/10 1.220 ± 0.214; n=56 cells). Thus, identification of SAT-induced synaptic plasticity concentrated in SST-Calb2 neurons enabled us to identify unique activity features of SST subtypes that showed strong predictive value for response plasticity during learning.

Intersectional genetic methods for cell-type identification are experimentally cumbersome and can suffer from a high false-positive rate due to low levels of marker gene expression in other cell classes, particularly true for the Calb2 gene (*25*). Label-free cell clustering using features of neural activity would be a major advance in deciphering cell identity from large-scale imaging or recording experiments. Thus, we asked whether this classifier might enable us to identify putative SST-Calb2 neurons from the larger unlabeled dataset, that is, neurons from SST-Cre transgenic mice (n=98 cells in 10 mice). Neurons with transgenic expression of GCaMP6f were typically brighter than virally labeled cells, complicating application of a fixed-parameter classifier that would not generalize between datasets, a potential problem that was addressed using clustering-informed classification. Again, classification was carried out using data only from the pretraining ACC period.

Similar to results from the labeled SST-Calb2 dataset in Fig. 4D, cluster analysis segregated SST neurons into multiple groups, one of which was assigned as putative SST-Calb2 based upon similarity of their response properties to the SST-Calb2 cells (Fig. 4F, J). Again, we used this grouping to ask whether these putative SST-Calb2 neurons would selectively show response suppression during learning. Classification of SST-Calb2 neurons from the ACC period accurately predicted response suppression during SAT, where this group showed a highly significant, 40% response suppression by the end of SAT, similar to results from Fig. 3E and Fig. 4D (putative SST-Calb2 fold-change versus ACC4-6 for SAT1/2 0.900 ± 0.053; SAT5/6 0.728 ± 0.078; SAT9/10 0.572 ± 0.074, n=30 cells). In comparison, SST neurons in the SST-O combined clusters showed no significant change during SAT (putative SST-O fold-change versus ACC4-6 for SAT1/2 0.924±0.056; SAT5/6 0.981±0.085; SAT9/10 1.013±0.095, n=68 cells).

Pseudotraining did not depress stimulus-evoked SST activity in superficial layers of S1; indeed, responses were increased in at least a subset of neurons (fig. S6). To determine whether putative SST-Calb2 neurons were modulated by pseudotraining, we applied the classifier to ACC (pretraining) data from this separate group of animals. This analysis revealed similar cell clusters (fig. S20), one of which we assigned as SST-Calb2 based upon response features. However, pseudotraining did not differentiate sensory-evoked responses in SST-Calb2 vs. SST-O neurons (fig. S20). These data suggest that plasticity in SST-Calb2 neurons is selectively initiated by predictive sensory-reward information but not when sensory inputs are unrelated to reward outcome.

## Discussion

Developing a census of molecularly- and functionally-defined neuronal cell types is an important step in understanding and replicating the computational properties of the neocortex, a 6-layered anatomical structure that is highly conserved across the neocortex and also mammalian species (*26*). Although cortical lamina provide a useful structure to understand cortical information flow (*27*), the diversity of neuronal subtypes and their connectivity even within a layer has complicated this interpretation (*24*, *28*). Transcriptomic studies indicate that there may be hundreds of distinct cell types in the neocortex, with substantial diversity even within defined subclasses of excitatory and inhibitory neurons (*13*). The correspondence of this transcriptomic diversity with neural activity and function remains poorly understood, particularly for cortical interneurons.

Although functional diversity in neocortical SST neurons has been suggested by recent studies (*8*, *18*, *29*), linking this diversity to molecularly-defined subclasses using transcriptomic approaches has been elusive (*17*). Our *in vivo* imaging and anatomical analysis indicate that SST-Calb2 Martinotti neurons in superficial layers of sensory cortex are selectively modified by stimulus-reward association during learning but not in pseudotraining, when whisker stimuli have no predictive value in water delivery. Thus, L2/3 SST-Calb2 neurons lie in a distinct node of the cortical circuit responsive to contingency information and are likely to be distinguished by their synaptic connectivity and/or selective responses to neuromodulators. Furthermore, chemogenetic inhibition of SST neurons was sufficient to initiate a loss of associated PSD95 synaptic sites onto *any* SST neuron in superficial layers, suggesting that the SAT-dependent effect does not result from a cell-type specific transcriptional program in SST-Calb2 neurons but may be directly related to reduced activity of this subclass during training.

Because SST activity was suppressed prior to a reduction in PSD95 puncta size during SAT, our data suggest that decreased synaptic PSD95 specifically in SST-Calb2 neurons may result from suppression of sensory-evoked activity in these cells. It will be of great interest identify the circuits involved in SST-Calb2 activity suppression during learning. Because SST neurons receive strong inhibition from GABAergic VIP neurons (*30*), changes in VIP activity during stimulus-reward association could be an early step in the SAT-dependent regulation of SST output.

Our experiments purposefully monitored SST activity outside of the training context, enabling us to avoid confounds from task-related activity (*31*) to focus on sensory-evoked responses. Because we also observed marked synaptic changes onto SST-Calb2 neurons that were preserved in fixed tissue, changes in the stimulus-evoked responses of SST neurons are likely to be present during training and can dynamically influence sensory processing.

Activity-based cell-type classification has been of great interest to neuroscientists, as it can provide structure to *in vivo* recordings that shed light on neural computation (*32*, *33*). The increasing number of molecularly-defined neural subtypes in the brain has complicated these efforts. Our anatomical results and cluster-based classification data validate transcriptomic and anatomical distinctions between SST subtypes, indicating that these neurons play selective roles in cortical reorganization during learning. Importantly, we employed a clustering-informed classification method that could adapt to variation in Ca^++^ signals collected from different genetic methods and experimental set-ups. This cluster-based classification approach provides a robust and easily-adopted method for label-free, cell-type identification from *in vivo* recordings, an important step in understanding the algorithms that underlie information processing in the brain.

## Acknowledgements

Special thanks to Molly Kinstle for immunolabeling and histological analysis, Alex Kireeff for technical assistance and cage design, Rachel Swindell for behavioral analysis code, Joanne Steinmiller, Rachel Bouchard, and Lyndsay Krut for expert animal care. Thanks also to members of the Barth lab, Aric Agmon, Eric Yttri, and Caroline Runyan for critical comments and suggestions throughout the project.

## Author contributions

Conceptualization: MBM, XM, MZ, ALB Methodology: MZ, MBM, XM Investigation: MBM, MZ, XM Visualization: MZ, MBM, XM Funding acquisition: ALB Project administration: ALB Supervision: ALB, MZ, MBM Writing – original draft: ALB Writing – review & editing: MBM, MZ, XM, ALB

## Data and materials availability

All code used for analyses can be found at: https://github.com/barthlab/Long-lasting-subtype-specific-regulation-of-somatostatin-interneurons-during-sensory-learning. All data used for graphing can be found at https://doi.org/10.5061/dryad.qbzkh18st. All other data will be provided by the authors at the reader’s request.

## Materials and Methods

### Animals

For GCaMP6f imaging in somatostatin neurons, we crossed Sst-IRES-Cre mice (Jackson #013044) to Ai148(TIT2L-GC6f-ICL-tTA2)-D mice (Jackson #030328). For GCaMP6f imaging in calretinin-expressing somatostatin neurons, we crossed Sst-IRES-Flp mice (Jackson #028579) to Cr-IRES-Cre (Calb2-IRES-Cre) mice (Jackson #010774). Juvenile to adult transgenic mice (1.5-6 mos of age) were used for cranial window surgery and virus injection. They recovered for 1-3 weeks before commencing 2P *in vivo* imaging. Sst-IRES-Cre mice (Jackson #013044) were used for fixed tissue analysis. Male and female mice were used for all experiments and approximately balanced across control and experimental datasets.

### Cranial window surgery

Surgery was done under isoflurane anesthesia (4% for induction, 1.5-2% for maintenance). Mice were put on a heat pad with a temperature control system (FHC 40-90-8D) to maintain body temperature. Eyes were covered with Puralube Vet Ointment to prevent drying. Fur was removed with Nair, and the skin was cleaned with povidone and then incised expose the skull. The skull was scraped with a dental blade (Salvin 6900) to remove the periosteum and abraid the surface for headpost attachment. On the left hemisphere, S1 coordinates (3.5 mm lateral, 1 mm posterior to bregma) and a 3 mm diameter circle centered at the coordinates were marked with a pen. A thin layer of tissue adhesive (3M VetBond) was applied to the skull, then a custom-made headpost was attached to the right hemisphere with cyanoacrylate glue and dental cement (Lang Dental, 1223PNK). With a dental drill (Dentsply, 780044), the skull was thinned along the 3 mm diameter circle. Thinned skull was removed by lifting a spot of the thinned region with forceps. Minor bleeding was stopped with saline-soaked gelfoam (Pfizer, 00009032301), and a glass window comprised of a 3 mm diameter glass (Warner Instruments, 64-0726) attached to a 4 mm diameter glass (Warner Instruments, 64-0724) by UV adhesive (Norland, 717106) was applied over the craniotomy. The window was sealed with 3M Vetbond and then cyanoacrylate glue. All exposed skull area except the window was covered with dental cement. A well surrounding the window was built with dental cement for microscopy using a water immersion lens. At the end of the surgery, ketoprofen (3 mg/kg) was injected subcutaneously, and the mouse was allowed to recover in a heated cage. Mice were given 1-3 weeks of recovery before imaging commenced.

### Stereotaxic injections

Male and female mice expressing Cre recombinase under the somatostatin promoter aged P50-P100 were induced into anesthesia with 4% isoflurane and administered a maintenance dose of ∼1.5% isoflurane throughout the duration of surgery. Using a dental drill, a burr hole was drilled preserving the final layer of skull to minimize damage to the cortex. Saline was washed over the skull to soften the injection zone prior to insertion of the glass pipette. Mice were injected with ∼80nL of AAV-PHP.eB-ZFN-hSyn-DIO-PSD95.FingR-Citrine-reg.WPRE into S1 (−3.5mm lateral, −1.25mm posterior relative to lambda) using a nanoject (Drummond Scientific). Post injection, sutures were used to close the scalp lesion. Following surgery, mice were single housed to reduce possible confounds of different socialization levels among mice compared across trained or naïve conditions.

For chemogenetic experiments designed to monitor changes in PSD95 puncta after suppressing activity, we coinjected pAAV8-hSyn-DIO-hM4Di-mCherry (Adgene #44362) with AAV-PHP.eB-ZFN-hSyn-DIO-PSD95.FingR-Citrine-reg.WPRE into S1BF using the stereotaxic injection procedures outlined above. Aliquots of Clozapine-N-Oxide (CNO; ApexBio) made up in DMSO (0.01mg/ul) were placed in the drinking water to approximate the dosage of 1mg/kg per day based on animal initial weight and estimated water consumption of 2 ml per day. To administer CNO, mice were placed in the training cage without any airpuff cue and were freely able to collect water containing CNO for five days. Like the acclimation period in other SAT training experiments, trials dispensed water at 80% probability.

For GCaMP6f imaging and calretinin labelling in SST neurons using viral methods, we injected AAV1-Ef1a-fDIO-GCaMP6f (Addgene #128315) mixed at a 1:1 ratio with AAV8-Ef1a-Con/Fon-mCherry (Addgene #137132) or pAAV8-hsyn-DIO-mCherry (Addgene #50459) in Sst-IRES-Flp x Cr-IRES-Cre (Calb2-IRES-Cre) mice using a Nanoject II (Drummond Scientific) directly before application of the cranial window implant. After craniotomy, a ∼0.6 µL of virus (∼1.84×10¹³ vg/mL) was injected into the barrel cortex (three sites across the cranial window, 0.3 mm below the pial surface). The virus expression period lasted 26-42 days prior to the commencement of imaging.

### Sensory association training (SAT)

Mice were trained to associate a multiwhisker stimulus with a delayed water reward in an automated training cage (*14*, *34*). Mice were single-housed in a home cage connected to a freely-accessible chamber with a water port and an airpuff delivery tube. Animals were not water deprived. During the cage acclimation period, animals could freely approach the lickport and initiate a trial through breaking an infrared beam. During the acclimation period, water delivery trials consisted of a delay period lasting 1.2-1.8 seconds preceding water delivery where a water droplet (∼10 ul) was dispensed at 80% probability (i.e. 20% of nosepokes did not result in water delivery). During SAT, water delivery was preceded by a gentle airpuff (6 psi, 500 ms duration) to the right-side whiskers 1 second before water delivery. Anticipatory licking frequency was assessed during the 500 ms period following air puff stimulation and prior to water delivery. The sensory cue was fully predictive; i.e. all airpuff stimuli were followed by water. During SAT, 80% of trials consistent of the predictive airpuff followed by the water reward. The remaining 20% of trials had no stimulus and no water reward (blank trials). There was a 2s timeout between trials, where nosepokes would not trigger water delivery.

During pseudotraining, the airpuff stimulus randomly preceded either water or blank trials, so that airpuff had no predictive value for water delivery. During the acclimation period, water was delivered with a 50% probability to match the probability used for pseudotraining. During pseudotraining, airpuff was delivered in 80% of the trials but water followed the stimulus for only half the trials while the other half of stimulus trials were followed by no water delivery (fig. S3-4; (*14*). To further decouple the stimulus from the reward, water was delivered without a preceding airpuff for half of the remaining non-stimulus trials. Therefore, the airpuff stimulus and water reward were entirely decoupled during pseudotraining. Animal performance was calculated as described for SAT.

Both SAT and pseudotrained animals undergoing 2P imaging experienced 6 days of acclimation and then 10 days of training. For each animal, total number of trials (water + blank trials) and anticipatory lick frequencies (licks occurring in a 300 ms window right before water delivery; see (*34*)) were calculated for every 4-hour bin using a custom MATLAB code. Any 4-hour bin with fewer than 10 trials was removed from the averaged data, since lick frequency on blank trials could not be accurately assessed from 1-2 trials. Performance was calculated by taking the difference between anticipatory licking frequencies (Hz) during stimulus versus blank trials (licking_stimulus_-licking_blank_). Absolute differences in calculated lick frequency between stimulus and blank trials for the last 20% of trials on a given day were compared using a Wilcoxin signed-rank test.

### 2P *in vivo* imaging

All training sessions were conducted within the automated homecage training system, ensuring a consistent and controlled environment for behavioral learning tasks. For 2P imaging experiments, mice were removed from their homecage environment for brief periods of 1 hour per day, typically around noon. No lickport was present during training, and the number of stimulus trials was kept to a minimum (∼15) to prevent extinction. No difference in performance after imaging sessions was observed, indicating that this brief exposure did not alter the learned association (*35*). Mice were removed from the training cage around noon each day, briefly anaesthetized with volatile isofluorane (4% for ∼20 s) to headfix the animal under the microscope, and then allowed to recover for 3-5 minutes before imaging. Animals were awake and ambulatory on the wheel before imaging began. Imaging was carried out with 2P microscope (Femto2D Galvo), equipped with a Mai Tai laser MTEV HP 1040S (Spectra-Physics), a 4x air objective lens (Olympus UPLFLN 4X NA 0.13), and a 40x water objective lens (Olympus LUMPLFLN 40XW NA 0.8). Images were acquired with MES software v.6.1.4306 (Femtonics).

Blood vessel morphology in 4x brightfield was used to find the same imaging field of view (FOV) as the previous session. The pial surface (z=0) was defined as the plane right below the dura mater which looks like a textured membrane in 40x brightfield. In 40x 2P mode, the x, y, z positions of the neurons were aligned to match the previous session image. A 950 nm excitation was used to image GCaMP6f signals, and emission fluorescence was detected with photomultiplier tube (PMT; Hamamatsu H11706P-40). Laser power and PMT voltage were kept constant for each animal across its imaging sessions. Images were acquired at 5.11 Hz with ∼270 µm x 300 µm FOV and 0.7 µm/pixel resolution. Imaging depth was ∼200 µm below pia (L2/3), and 1-2 FOVs were imaged per mouse.

For each day, approximately 3-5 minutes after head fixation, 1-2, 10-minute imaging sessions were carried out, with a 1-minute break in between. At the beginning of each imaging session, spontaneous activity prior to sensory stimuli was recorded over a 100s window. Responses to either a vertical airpuff (500 ms duration, 6 psi) or blank (solenoid click) delivered by Arduino every 20 s (0.05 Hz) to the right-side whiskers during each session were obtained. Airpuff and blank stimuli had an equal probability of occurring and were randomly interleaved. We collected another 100s of spontaneous activity following stimulation. Following the imaging sessions, mice were promptly returned to their homecage training environment to minimize disruption to their daily routine and ensure the stability of their behavioral training regimen. Behavioral data from one mouse in the SAT SST-Cre x Ai32 dataset and 3 mice in the SAT SST-Flp x Calb2-Cre dataset were corrupted and unable to be analyzed. Animal participation in training could be deduced through tracking water consumption each day.

After animal training and when all imaging sessions were completed, the head bracket and window were removed, and the imaging site was marked by marking the site with a glass micropipette containing methylene blue dye. Brains were fixed in 4% paraformaldehyde and sectioned either coronally or flattened and cut tangentially to confirm the imaging site location.

### *In vivo* 2P analyses

An imaging file containing all imaging sessions (∼96000 frames) was aligned and segmented with Suite2P (*36*). The output from Suite2P included all possible segments. ROIs were then manually selected from all segments based on morphology and fluorescence traces calculated by Suite2P. Individual regions of interest (ROIs) (neurons) were tracked across each imaging day, and neurons that could not be tracked across all days were discarded from the analysis.

Image movement was assessed by calculating shifts in aligned pixels across frames, extracted from Suite2P. We established that any frame that shifted more than 20 pixels in either the X or Y direction within the larger FOV was considered a shifted frame. One or two continuously shifted frames were interpolated with the average value of the previous and the next unshifted frames (both fluorescence signal and pixel shift). When >3 consecutive frames were shifted within a single trial, the trial was then removed.

Raw fluorescence was extracted for each segmented ROI, and fluorescence signals were neuropil-corrected (F_corrected_ =FROI – 0.7*F_neuropil_) to remove a contribution from SST neurons in other layers (*37*). Baseline fluorescence (F_0_) was calculated by averaging the neuropil-corrected signal (F_corrected_) within a 1s time window preceding the stimulus onset of individual trials. The change in fluorescence relative to baseline, ΔF/F_0_, was computed for each trial using F_corrected_ - F_0_/F_0_. Individual neuropil-corrected ROIs were designated as neurons.

Responsive neurons were determined on each day, including both acclimation and training days using the following criteria. Peak response was defined by the maximum amplitude during the 1s window after the airpuff onset (5 image frames). Neurons were scored as responsive in a given trial if the peak ΔF/F_0_ >2 SD, where SD was calculated using the 1s window before the airpuff onset. The daily stimulus-evoked activity of each responsive neuron was calculated by averaging the cell response across all responsive stimulus trials within each imaging day. The peak response from ACC4-6 was used to normalize responses from the SAT period since neural activity during the first three imaging days showed greater variability than subsequent days of imaging in the pretraining period.

### Classification of calretinin (Calb2+) neurons

To identify Calb2 neurons from *in vivo* imaging FOVs in SST-Flp x Calb2-Cre mice, an intensity matrix for each image plane was initially extracted from MES and subsequently transformed into a two-channel image featuring green (GCaMP6f) and red (mCherry) channels. The cell bodies of SST neurons were manually traced based on GCaMP6f expression using ImageJ on both the initial (ACC1) imaging day within each imaging plane. Following this, the average pixel intensity of each corresponding region of interest in the mCherry channel was calculated. A cell was categorized as Calb2 positive if its average pixel intensity exceeded 200 on the initial imaging day (fig. S13).

### SST activity feature extraction

Activity features of neurons were extracted from GCaMP6f fluorescence signals from 10-minute long sessions as described above, only from ACC4-6 imaging sessions. For each cell in a session, the global baseline (for the entire session, F₀) was computed by averaging the fluorescence from the concatenation of all 1-second intervals immediately preceding stimulus or blank onset. The fluorescence change was calculated as F-F_0_/F_0_ (i.e. ΔF/F_0_). To remove long-term trends, a 5-second kernel median filter was applied to the normalized fluorescence signal. The median absolute deviation (MAD) of the detrended ΔF/F_0_ was calculated as a measure of noise.

Individual sessions were divided into two distinct periods: the evoked response period (from −3s to +4s relative to stimulus/blank trial onset) and the spontaneous activity period (the remaining portion of the trace; note that this assumes stimulus-related activity has concluded after 4s). For the spontaneous activity period, peaks were identified using SciPy’s *find_peaks* function (version 1.11.1). From these spontaneous events, various features were computed, including the mean and median of the highest peak, prominence, inter-event interval, peak duration, relative height, and frequency.

For the evoked response period, each trial was further segmented into four intervals: *pre* (−3s to 0s), *evoked* (0s to 1s), *late* (1s to 4s), and *post* (0s to 3s) relative to trial onset. The peak value of detrended fluorescence (ΔF/F₀) was calculated for each segment. The average peak values were then computed for stimulus trials, blank trials, and then all trials (stimulus+blank) as features. Additionally, response probabilities were determined by calculating 2 ratios based on calculation of a “significant response,” where peak fluorescence exceeded a specified threshold of MAD values (fig. S16). These ratios were 1) significant response trials versus all trials and 2) significant stimulus response trials versus all stimulus trials. Due to broad variation in the distribution of peak response amplitude and background fluctuations in fluorescence across individual neurons in our dataset, the “significant response” feature was assessed using multiple thresholds (2x, 3x, 5x, 10x, and 20x MAD values) for detection of peak fluorescence of the evoked period. Extracted features used for classification are described in Tables 1-3.

### UMAP clustering and clustering-informed heuristic classification

Fixed-parameter classifiers can fail to perform robustly across varied experimental conditions. Differences in mouse lines, calcium imaging environments, and equipment can significantly alter fluorescence distributions, introducing hard-to-quantify shifts in the distribution of the data. As a result, these static classifiers are prone to overfitting (particularly with small datasets) and often fail to capture the underlying structure of heterogeneous datasets, contributing to non-reproducibility across studies. Thus, we chose to use clustering-informed heuristic classification to investigate the intrinsic structure of SST activity profiles present in both transgenic and virally-expressed GCaMP6f signals, from pretraining data (ACC4-6).

Uniform Manifold Approximation and Projection (UMAP) dimensionality reduction (with n_neighbors set to 8; version 0.5.3) was applied to reduce the dimensionality of all ***Evoked Response Peak Features*** and all ***Evoked Response Probability Features*** (Table S2, S3; Fig. 4B, I; Fig. S18B). Features were averaged across recordings from ACC4-6, and then z-score normalized to ensure equal feature contributions.

After reducing the feature space to two dimensions, spectral clustering (with n_clusters set to 3, reflecting the three apparent clusters in the UMAP plot) was used to classify cells into distinct clusters. The setting that combined ***Evoked Response Peak Features*** and ***Evoked Response Probability Features*** generalized well across both labeled and unlabeled datasets. Because incorporating features from ***Spontaneous Activity*** (in a 100s period prior to stimulus delivery) generally degraded performance of the classifier (Fig. S17B), spontaneous activity features were not used for classification. Further details are available in the accompanying code: https://github.com/barthlab/Long-lasting-subtype-specific-regulation-of-somatostatin-interneurons-during-sensory-learning

### Evaluation of classifier accuracy and robustness

To validate the robustness and reproducibility of the clustering results, we calculated the Adjusted Rand Index (ARI) and Normalized Mutual Information (NMI) for each dataset. Based on the ACC4∼6 data, we generated 10 random clusterings (with *random_state* values from 0 to 9) and computed ARI and NMI scores for each pair of random seeds (Fig. S18). In the SST-Calb2 labeled dataset, the ARI and NMI were 0.67±0.02 and 0.76±0.01. Similarly, the ARI and NMI for the unlabeled SAT dataset were 0.87±0.02 and 0.91±0.01, demonstrating consistent clustering robustly emerged from different random clustering seeds. For the unlabeled dataset from pseudotrained animals, ARI and NMI values were 0.97±0.004 and 0.97±0.004. In terms of classification performance, the classifier applied on the SST-Calb2 labeled dataset achieved an F1-score of 0.57 when the outlier group was ignored, and 0.52 when the outlier group was classified as SST-O cells. The AUC-ROC was 0.68 without the outlier group and 0.64 when including it. The overall accuracy was 0.71 without the outlier group and 0.70 when it was included.

### Tissue collection for anatomical analyses

To control for the transduction time of PSD95.FingR, mice were sacrificed 14 days after viral injections at midday regardless of experimental condition. Mice were deeply anesthetized with a near lethal dose of isoflurane and transcardially perfused with 20mL of 1x PBS followed by 20mL of 4% paraformaldehyde in 1x PBS. Brains were carefully removed and post-fixed in 4% PFA overnight followed by transfer to 30% sucrose in 1x PBS. Approximately three days following sucrose immersion ∼50um thick free-floating sections were acquired using a freezing microtome (Leica Biosystems) and stored in PB. Sections typically underwent immunohistochemical staining within two days of slicing.

### Immunohistochemistry

Prior to staining, four alternating sections containing posterior barrel cortex were washed in 1x PBS (Boston BioProducts Inc) for five minutes over five cycles. Sections were shaken in a blocking solution containing 1x PBS, 10% goat serum (Sigma-Aldrich) and .3% triton X (Sigma-Aldrich) in MilliQ water for 2 hours. After blocking, a 1:500 dilution of rabbit α calretinin primary antibody (Swant #CR7697) was mixed with the block solution containing 5% goat serum instead of 10%. Sections were covered and placed on a rocker at 4°C overnight (20∼24 hours). After the primary, sections were washed in 1x PBS for 5 minutes over five cycles. Finally, sections were shaken in a 1:500 dilution of Far-red secondary antibody (CF640R α Rabbit) in 1x PBS for 2 hours followed by a final step of 1x PBS washes. Sections were immediately mounted in antifade mounting media containing DAPI (Vectashield) on Diamond White Glass charged slides (Globe Scientific).

### Confocal image acquisition

Fields of view (FOVs) were collected using an LSM 880 Axio Observer microscope (Carl Ziess). Using a 10x objective, FOVs were coarsely targeted over S1BF using barrels resolved by DAPI staining as a guide. Precise targeting of L2/3 FOVs under the 63x oil immersion objective lens (Plan-Apochromat, NA 1.40, oil) was done using the granular layer (start of L4) demarcated by DAPI staining as a lower bound of and the steep drop off in SST dendritic arborizations as an upper bound of L2/3 (bottom of L1). L4 was targeted by using an increase in DAPI signal as a marker and generally 400-500um from the pial surface while L5 was characterized by an increase in density of SST somas. Volumetric stacks using the 63x oil immersion objective lens set at a .9 zoom factor and 1.0 Airy disk unit was used to collect ∼100 1024 x1024 pixel images with a z-step size of .3um. This resulted in a 149.54 x149.54 x 29.7um image stack with the voxel dimensions 0.146 x 0.146 x 0.3um. PSD95.FingR-Citrine fluorescence was collected for later surface reconstruction analysis (excitation 514nm, emission: 517-561nm). For experiments where PSD95.FingR puncta were registered with Calb2 identity, the far-red channel (excitation 640nm; Emission 641-695) was also captured. 514nm laser power was adjusted (generally between 4-6%) for each FOV to prevent over-or under saturation of punctate PSD95.FingR pixel intensities. The gain was set to be between 700-720 arbitrary units across all animals.

### Digital reconstruction of fluorescent signal

Volumetric stacks containing PSD95.FingR-Citrine labeled excitatory synapses were analyzed using the image analysis software Imaris (version 8.4.1; Bitplane). The citrine channel was adjusted such that background signal was subtracted to resolve PSD95.FingR puncta. This fluorescent channel was then digitally reconstructed using a the Imaris watershedding algorithm (Surfaces macro; image segmentation). The watershed threshold for PSD95 puncta signal (without smoothing; expected size 0.5um) was adjusted to maximize coverage of fluorescence signal while minimizing reconstructing signal contained in the background. Fused objects (adjacent overlapping signal) were split using the quality filter feature built into the surface macro (0.45um). Surfaces were then filtered by voxel size (>3 voxels) to minimize noise captured in surface reconstructions. In cases where all PSD95 puncta from a FOV were included in the analysis, reconstructions resulting from somatic and nuclear citrine fluorescent signal were removed by filtering surfaces <0.15um from reconstructed somas (watershed threshold: smoothing 0.263 arbitrary units; no quality filter; voxel size: adjusted to include fully reconstructed somas).

### Confocal image analysis

Following digital reconstructions of fluorescent signal, characteristics of PSD95 puncta surfaces could be quantified and compared across conditions. Volumes of each reconstructed surface were obtained to estimate the size of putative excitatory synapses. These puncta were averaged together to obtain a mean size of excitatory puncta for a given animal or cell type. For large-scale FOV analyses (fig. S10), unlabeled puncta obtained from each animal were randomized and balanced across conditions.

To quantify the difference between levels of excitation in SST-Calb2 vs SST-O neurons, we registered puncta as belonging to somas that were positive or negative for immunohistochemically labeled calretinin (protein product of the Calb2 gene). Since SST neurons are relatively sparse, PSD95.FingR puncta on individual dendrites that emanate from an individual soma could be resolved. Following semi-automated reconstruction of the entire field of PSD95 surfaces, ∼200 puncta were manually selected along a stretch of dendrites emanating from a given soma of each cell type. Experimenters were blind to cell-type identity and condition during manual selection and later categorized puncta as belonging to Calb2 positive or negative neurons based on somatic calretinin expression. Both groups of SST neurons were captured in the same FOV and the image acquisition and reconstruction settings were held constant which enabled within-sample comparison between groups.

### Slice preparation

Animals were anesthetized with isoflurane briefly and decapitated between 11 am and 2 pm. 350μm thick off-coronal slices (one cut, 45° rostro-lateral) were prepared in ice-cold artificial cerebrospinal fluid (ACSF) composed of (mM): 119 NaCl, 2.5 KCl, 1 NaH2PO4, 26.2 NaHCO3, 11 glucose, 1.3 MgSO4, and 2.5 CaCl2 equilibrated with 95%/5% O2/CO2. Tissues were recovered at room temperature in cutting ACSF for 45 minutes to 1 hour before recording.

### General Electrophysiology

Recordings were performed in cutting ACSF in the presence of synaptic blockers (10µM NBQX, 50µM D-APV and 50µM PTX). Cortical SST neurons were targeted using an Olympus light microscope (BX51WI) and borosilicate glass electrodes (4-9 MΩ pipette resistance) filled with internal solution composed of (in mM): 125 potassium gluconate, 10 HEPES, 2 KCl, 0.5 EGTA, 4 Mg-ATP, 0.3 Na-GTP, and trace amounts of AlexaFluor 568 (pH 7.25-7.30, 290 mOsm) for morphological confirmation of cell identity. Electrophysiological data were acquired using Multiclamp 700B amplifier (Axon Instruments) and digitized with a National Instruments acquisition interface (National Instruments). Multiclamp and IgorPro 6.0 software (Wavemetrics) with 3kHz filtering and 10kHz digitization were used to collect data.

ChR2-expressing L2/3 SST neurons were targeted using reporter fluorescence. After breaking into the cells, SST neurons were voltage-clamped at −70mV for 3-5 minutes until the baseline membrane potential stabilized. Action potentials were evoked with depolarizing current steps (25, 50, 100, 150, 200, 250, 300pA, for 500ms duration, 3 sweeps each at 0.1Hz) in current clamp at −60mV. Cell identity was confirmed by both eYFP fluorescence, low-threshold spiking firing phenotype and light-evoked spikes. Only SST neurons with membrane potential ≤-40mV and a stable membrane potential baseline were included in our analysis. To calculate spiking frequency, depolarizations that exceed 0mV were counted as action potentials. The number of evoked action potentials was calculated by averaging 3 sweeps for each current injection amplitude. Resting membrane potential was obtained once cells had stabilized. Input resistance (MΩ) was calculated for each trial and averaged across all excitability experiment trials. Rheobase (pA) was determined as the minimum current required to elicit a single spike.

---

**Fig. S1:**
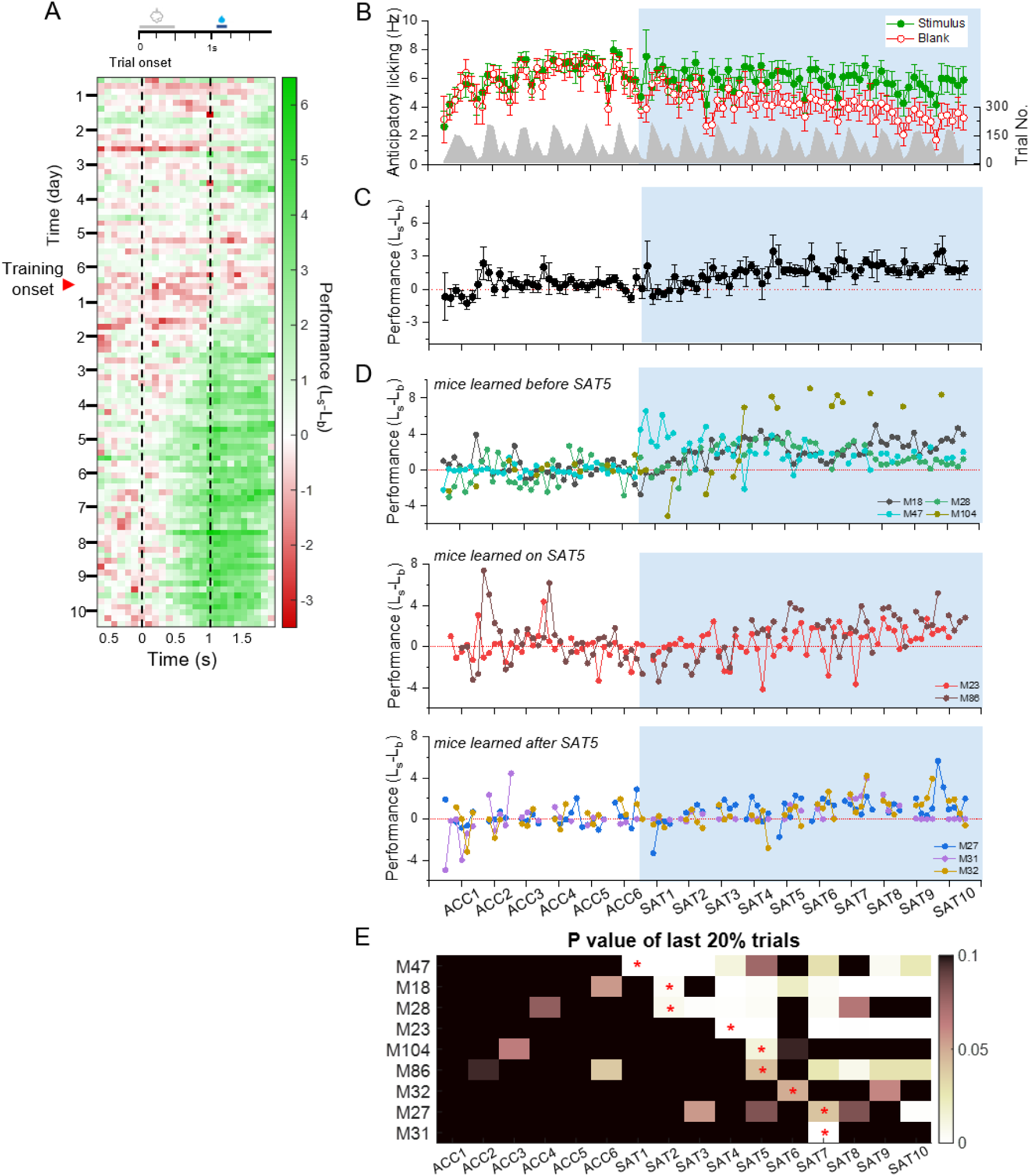
Anticipatory licking reflects learning during sensory association training (SAT) (**A**) Mean performance (Licking_stimulus_-Licking_blank_) averaged across 9 animals. (**B**) Mean anticipatory licking frequency averaged across 9 mice on stimulus (green) and blank (red) trials. (**C**) Mean performance averaged across 9 mice. (**D**) Top: Performance of individual mice learned before SAT5. Middle: same as top, but for mice learned on SAT5. Bottom: Same as top, but for mice learned after SAT5. (**E**) P values of anticipatory licking frequency of the last 20% of stimulus and blank trials for each animal. Asterisks indicate the learning day (see methods: criteria for learning).

**Fig. S2:**
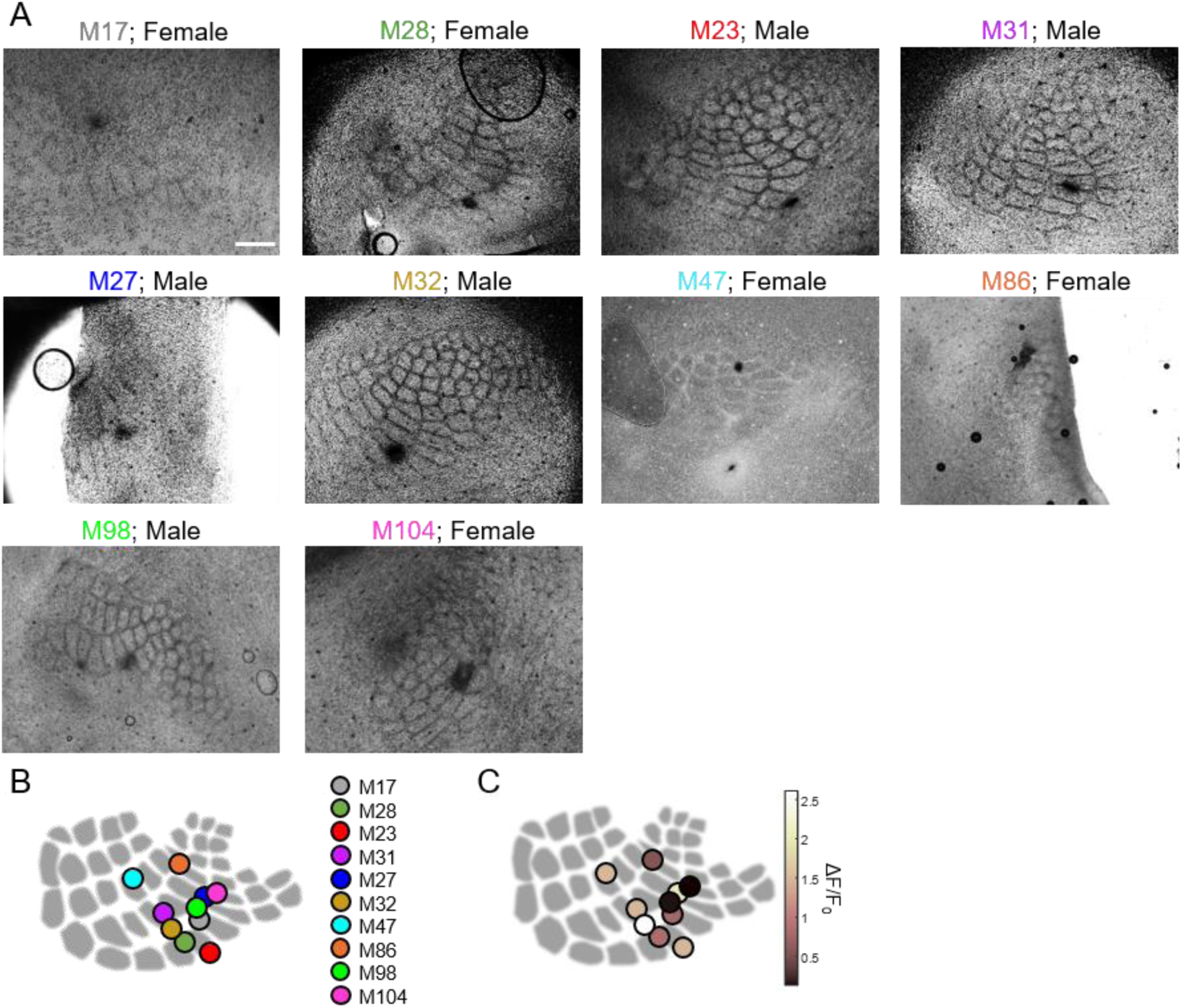
Post-hoc imaging location for SAT mice. (**A**) Post-hoc labeling of imaging site with a glass pipette filled with methyl blue dye. Scale=500 µm. (**B**) Schematic diagram of labeled imaging sites of all mice. Each dot represents a mouse. (**C**) Schematic diagram of labeled imaging sites of all mice color coded based on the mean peak response on ACC4-6.

**Fig. S3.**
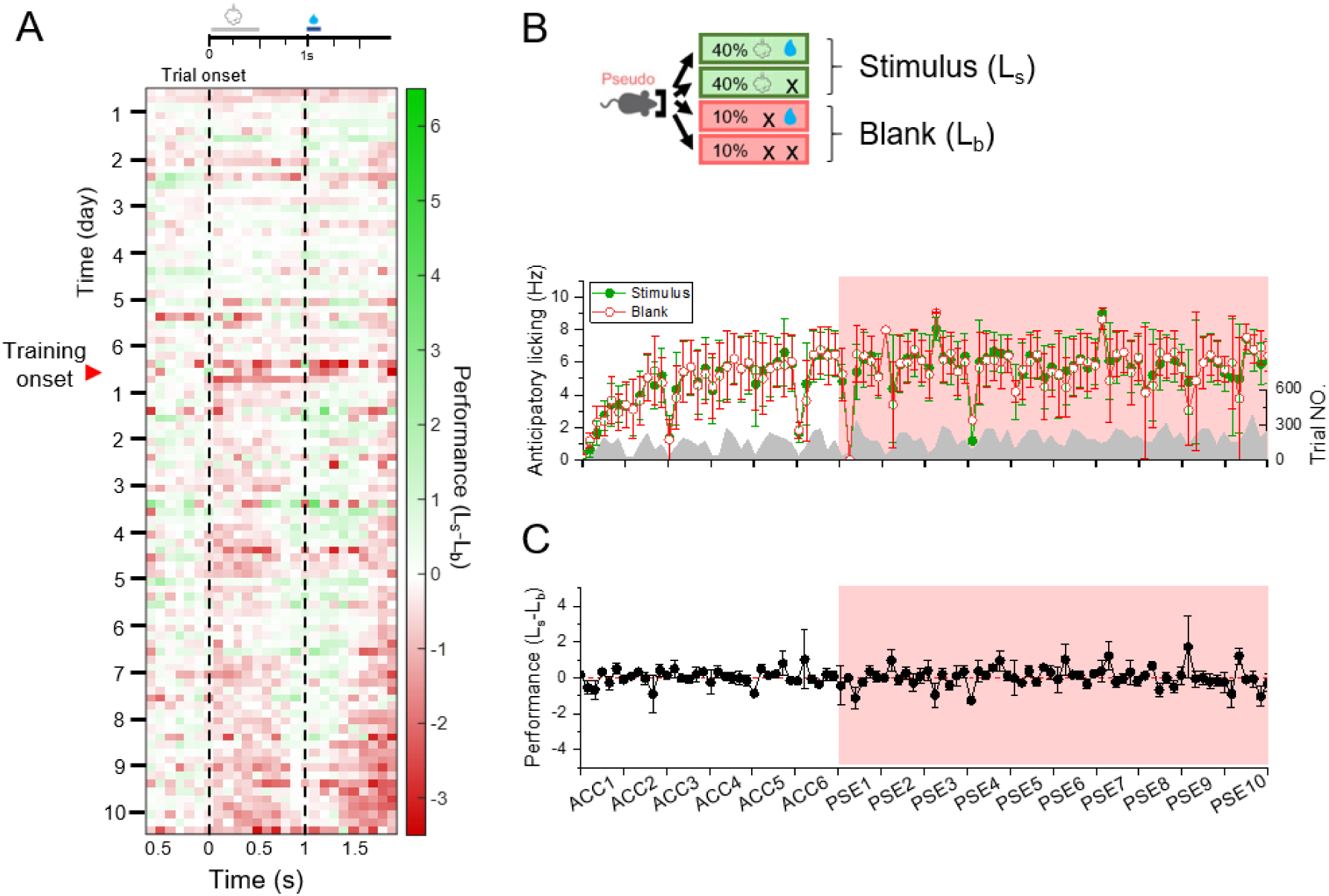
Stimulus-reward decoupling during pseudotraining (PSE) does not change anticipatory licking. (**A**) Mean performance (Licking_stimulus_-Licking_blank_) averaged across 10 animals. (**B**) Top: pseudotraining structure. Bottom: Mean anticipatory licking frequency averaged across 10 mice on stimulus (green) and blank (red) trials. (**C**) Mean performance averaged across 9 mice.

**Fig. S4:**
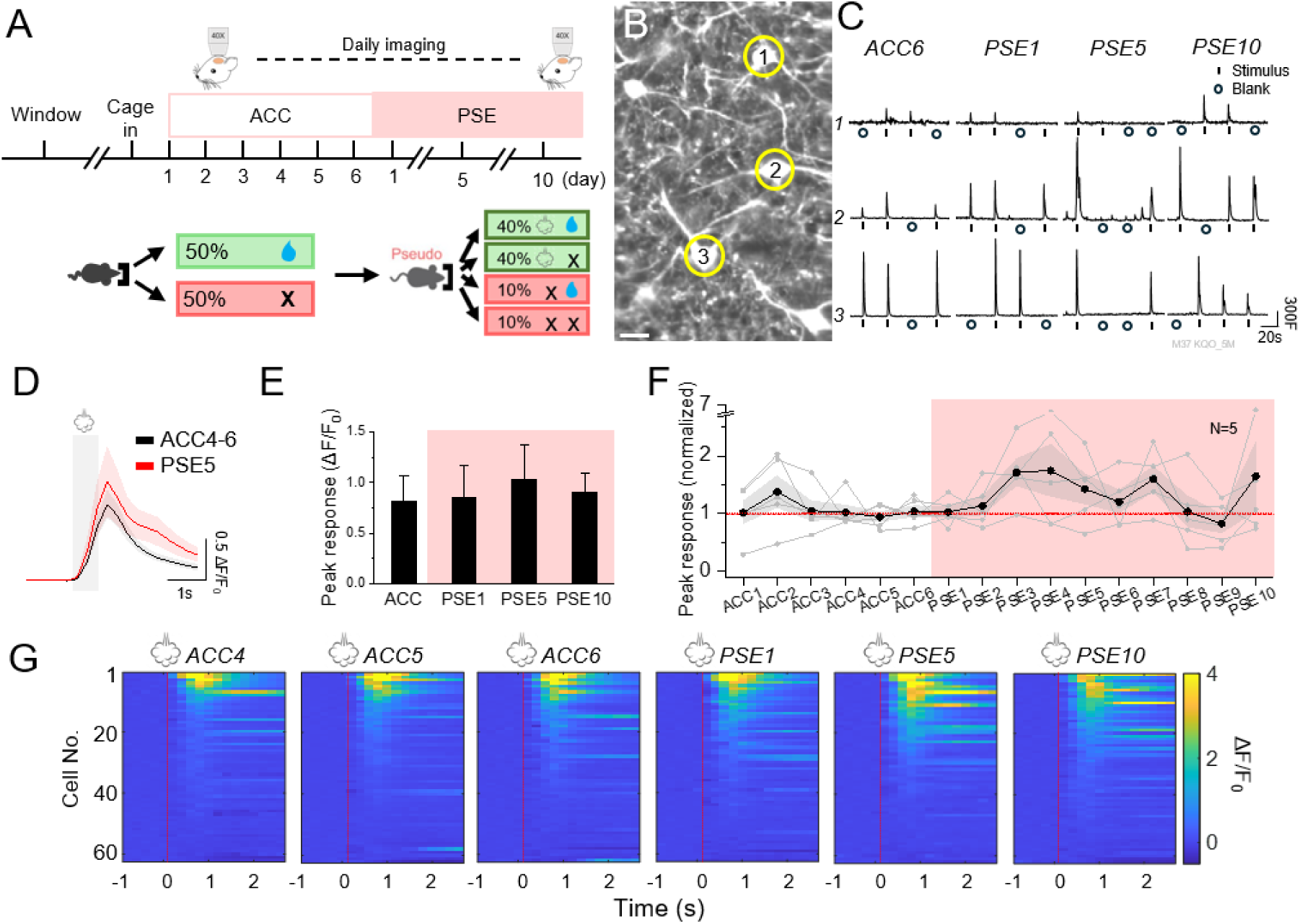
Pseudotraining enhances stimulus-evoked response in SST neurons. (**A**) Top: experimental timeline for longitudinal calcium imaging. Bottom: Structure of pseudo-training. (**B**) An example imaging FOV. Scale=20 µm. (**C**) Example traces of airpuff-evoked responses from the same example cells on ACC6, PSE1, PSE5, and PSE10. F (fluorescence) is reported in arbitrary units. (**D**) Averaged trace of the airpuff-evoked response on ACC4-6 and PSE5. Trace was averaged across 5 mice, 62 cells. Mean ± SEM of shown in the figure. Grey shade indicate the airpuff period. (**E**) Peak evoked response on ACC4-6, PSE1, PSE5, and PSE10, averaged by animal. Paired t-test with Bonferroni correction, comparing ACC4-6 with training days, p=0.99, 0.99, and 0.99 for PSE1, PSE5, and PSE10, respectively. (**F**) Peak airpuff-evoked response across the acclimation and training period, averaged across mice and normalized to ACC4-6 (black; n=5 mice). Mean ± SEM of shown in the figure. Grey lines indicate mean cell response for individual mice. One-way repeated measures ANOVA, p=0.074. (**G**) Heatmaps of the evoked response traces of all cells rank ordered within individual imaging days. Red dotted lines indicate airpuff onset.

**Fig. S5:**
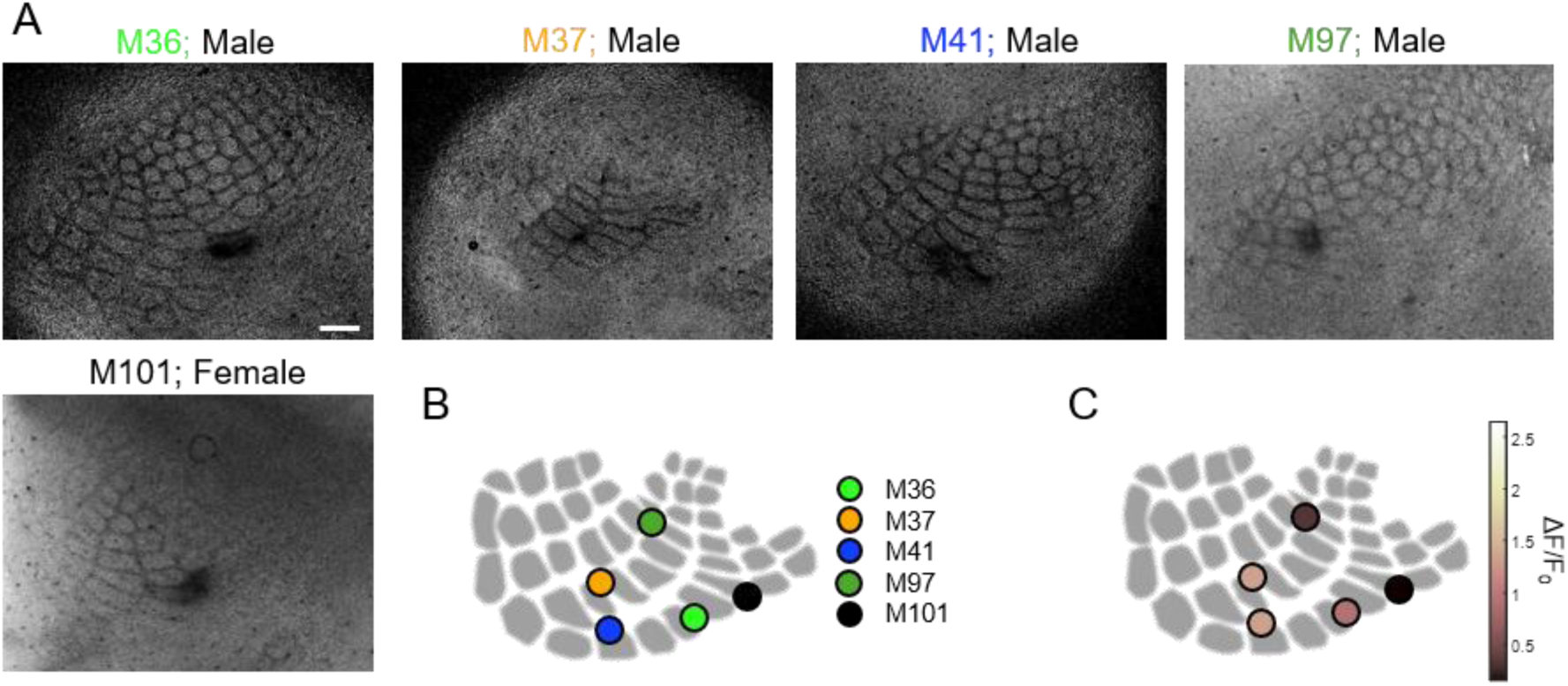
Post-hoc imaging location for PSE mice. (**A**) Post-hoc labeling of imaging site with a glass pipette filled with methyl blue dye. Scale=500 µm. (**B**) Schematic diagram of labeled imaging sites of all mice. Each dot represents a mouse. (**C**) Schematic diagram of labeled imaging sites of all mice color coded based on the mean peak response on ACC4-6.

**Fig. S6.**
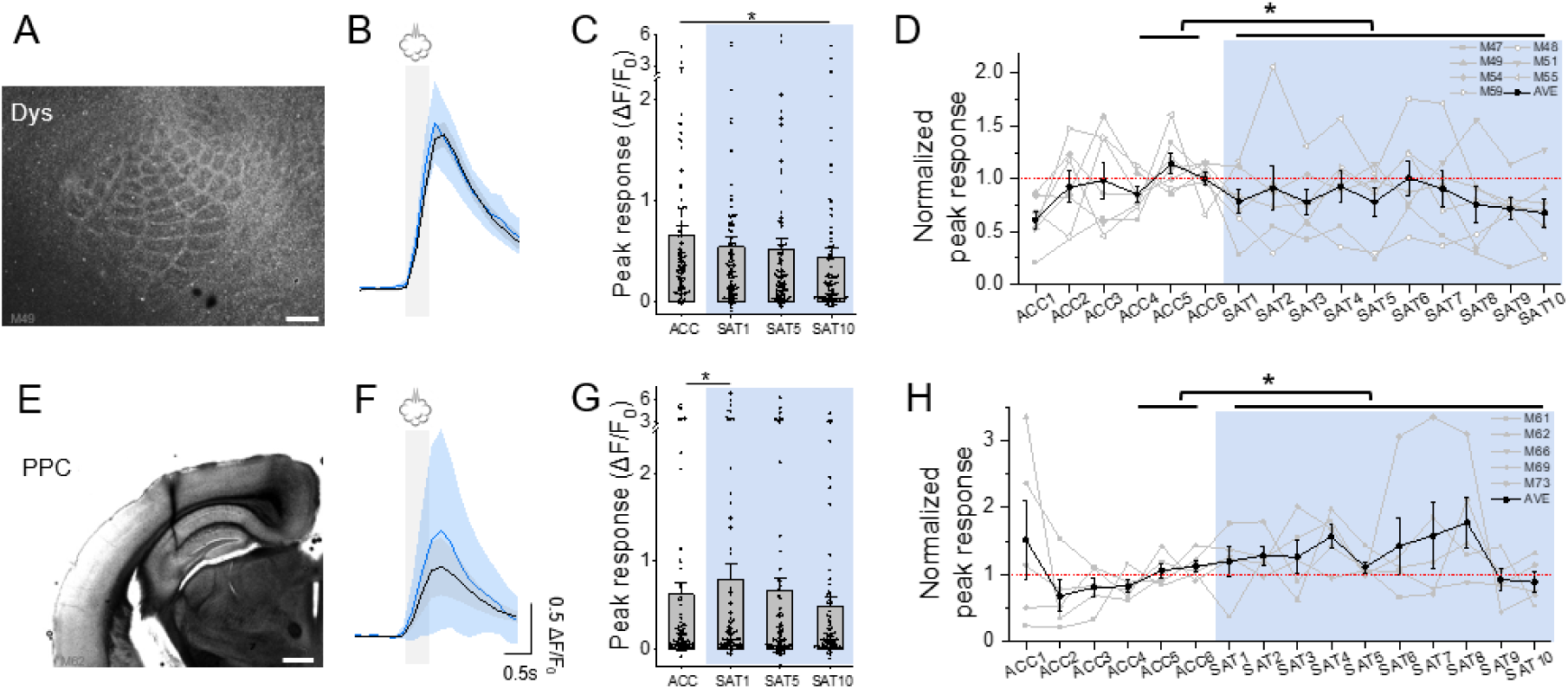
SST response plasticity outside of S1 barrel cortex. (**A**) Example imaging location in dysgranular zone. Scale bar = 500 µm. (**B**) Averaged trace of the airpuff-evoked response during acclimation 4-6 (ACC4-6) and SAT5. Trace was averaged across 7 mice. Mean ± SEM of shown in the figure. Grey shade indicate the airpuff period. (**C**) Peak evoked response on ACC4-6, SAT1, SAT5, and SAT10. Paired t-test with Bonferroni correction, comparing ACC4-6 with training days, p=0.17, 0.25, and 0.0076 for SAT1, SAT5, and SAT10, respectively. (**D**) Peak airpuff-evoked response across the acclimation and training period, averaged across mice and normalized to ACC4-6. One-way repeated measures ANOVA, p=0.0076. (**E**) Example imaging location in posterior parietal cortex. Scale bar = 0.2 mm. (**F**) Averaged trace of the airpuff-evoked response during acclimation 4-6 (ACC4-6) and SAT5. Trace was averaged across 5 mice. (**G**) Peak evoked response on ACC4-6, SAT1, SAT5, and SAT10. Paired t-test with Bonferroni correction, comparing ACC4-6 with training days, p=0.013, 0.99, and 0.17 for SAT1, SAT5, and SAT10, respectively. (**H**) Peak airpuff-evoked response across the acclimation and training period, averaged across mice and normalized to ACC4-6. One-way repeated measures ANOVA, p=0.013.

**Fig. S7.**
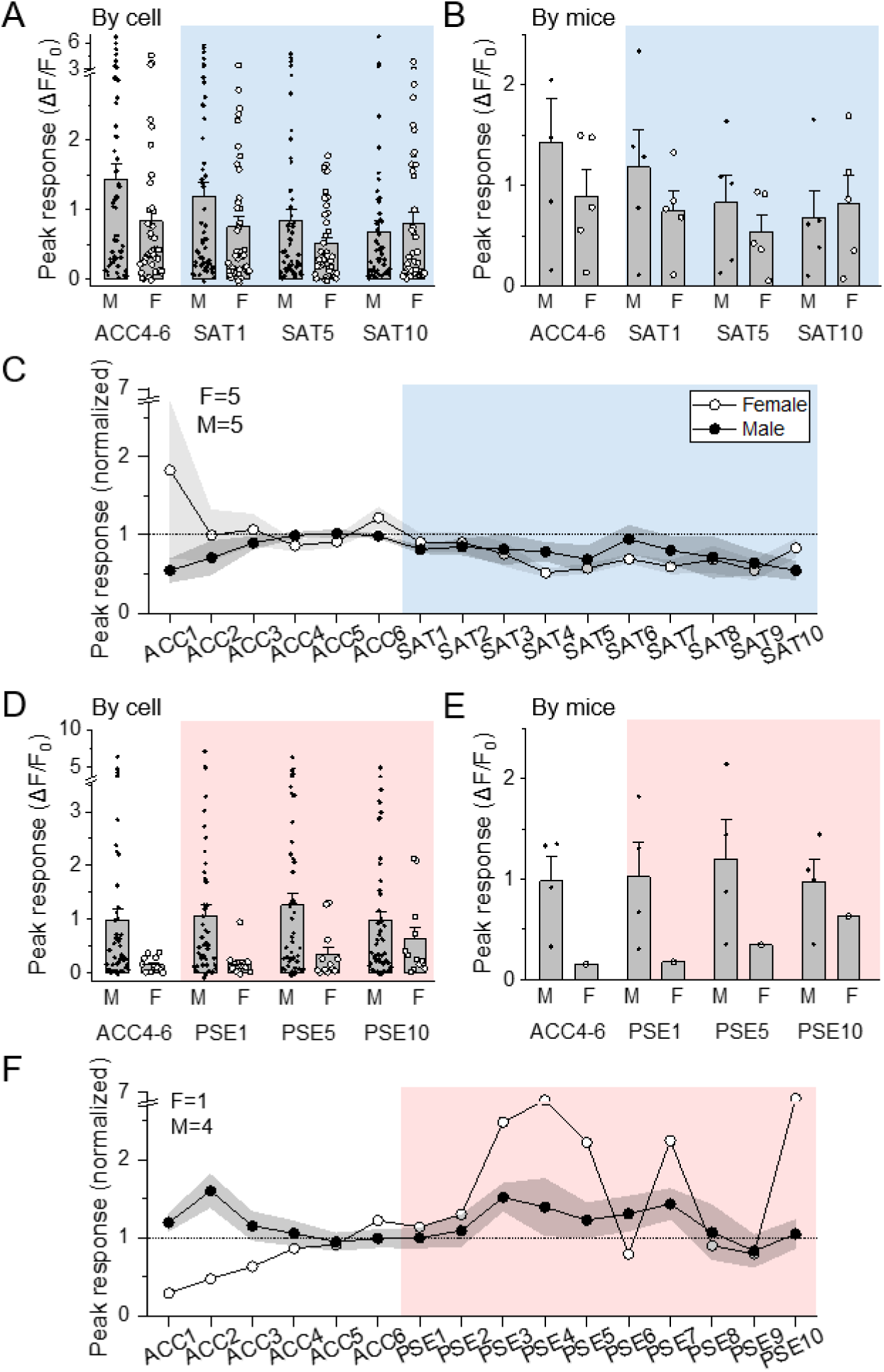
SST response plasticity in males and female mice. (**A**) Peak evoked response on ACC4-6, SAT1, SAT5, and SAT10 by sex, cell average. M: n=56, F: n=42. (**B**) Peak evoked response on ACC4-6, SAT1, SAT5, and SAT10 by sex, animal average. M: N=5, F: N=5. (**C**) Peak airpuff-evoked response across the acclimation and training period, averaged across mice, grouped by sex, and normalized to ACC4-6. Mean ± SEM of shown in the figure. (**D**) Peak evoked response on ACC4-6, PSE1, PSE5, and PSE10 by sex, cell average. M: n=50, F: n=12. (**E**) Peak evoked response on ACC4-6,, PSE1, PSE5, and PSE10 by sex, animal average. M: N=4, F: N=1. (**F**) Peak airpuff-evoked response across the acclimation and pseudotraining period, averaged across mice, grouped by sex, and normalized to ACC4-6. Mean ± SEM of shown in the figure.

**Fig. S8.**
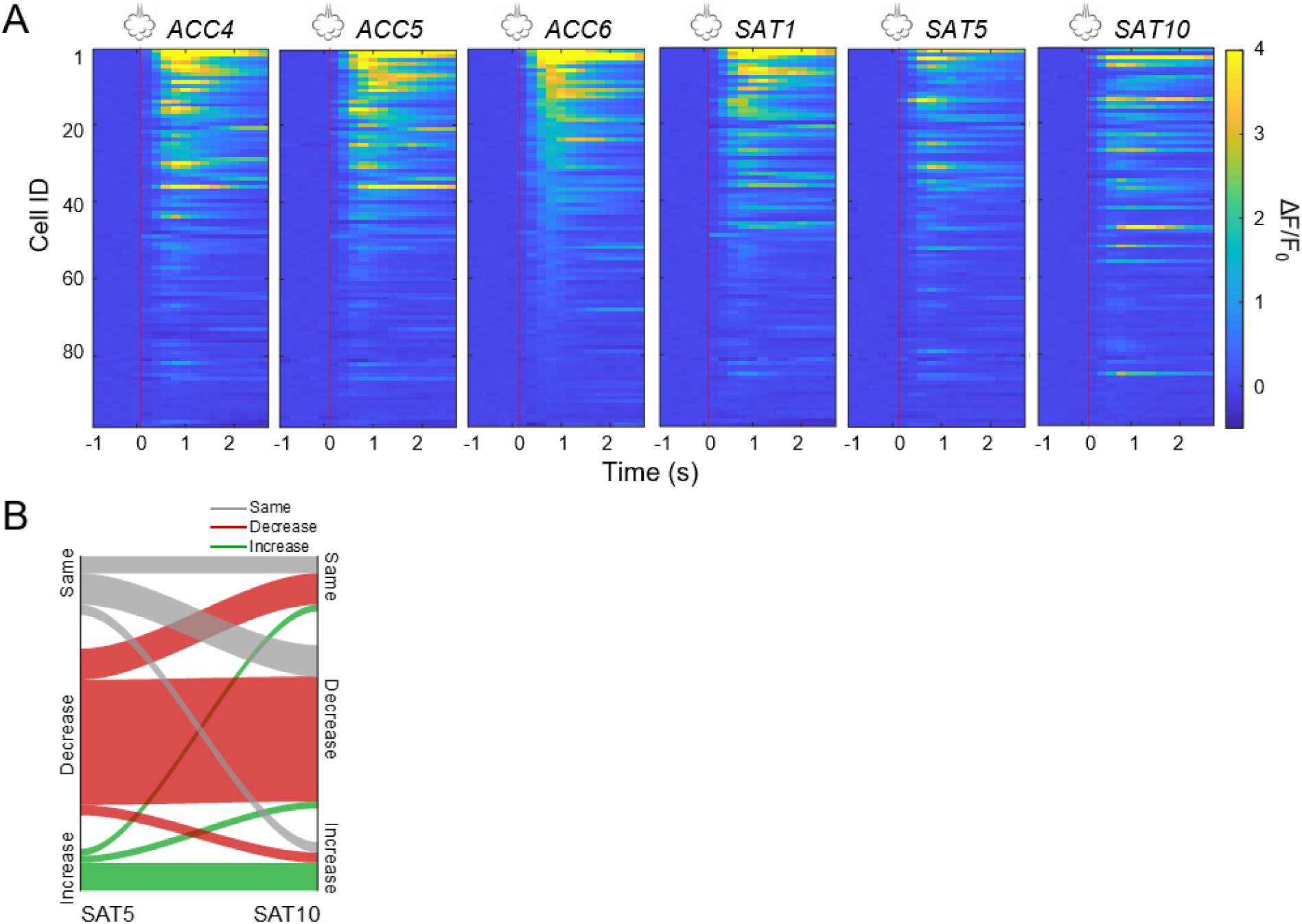
Tracking response plasticity in individual SST neurons during SAT. (**A**) Evoked response traces of all cells rank ordered by peak response from ACC6, where cell identity is maintained along the y-axis across days. Red dotted lines indicate airpuff onset. (**B**) Sankey plot showing changes in neural activity across sessions on SAT5 (left) to SAT10 (right). Categories are Same (gray; SAT5 n=17 and SAT10 n=16), Decrease (red; SAT5 n=48 and SAT10 n=47), and Increase (green; SAT5 n=12 and SAT10 n=14). Line thickness indicates the proportion of neurons transitioning between states on these two imaging days.

**Fig S9.**
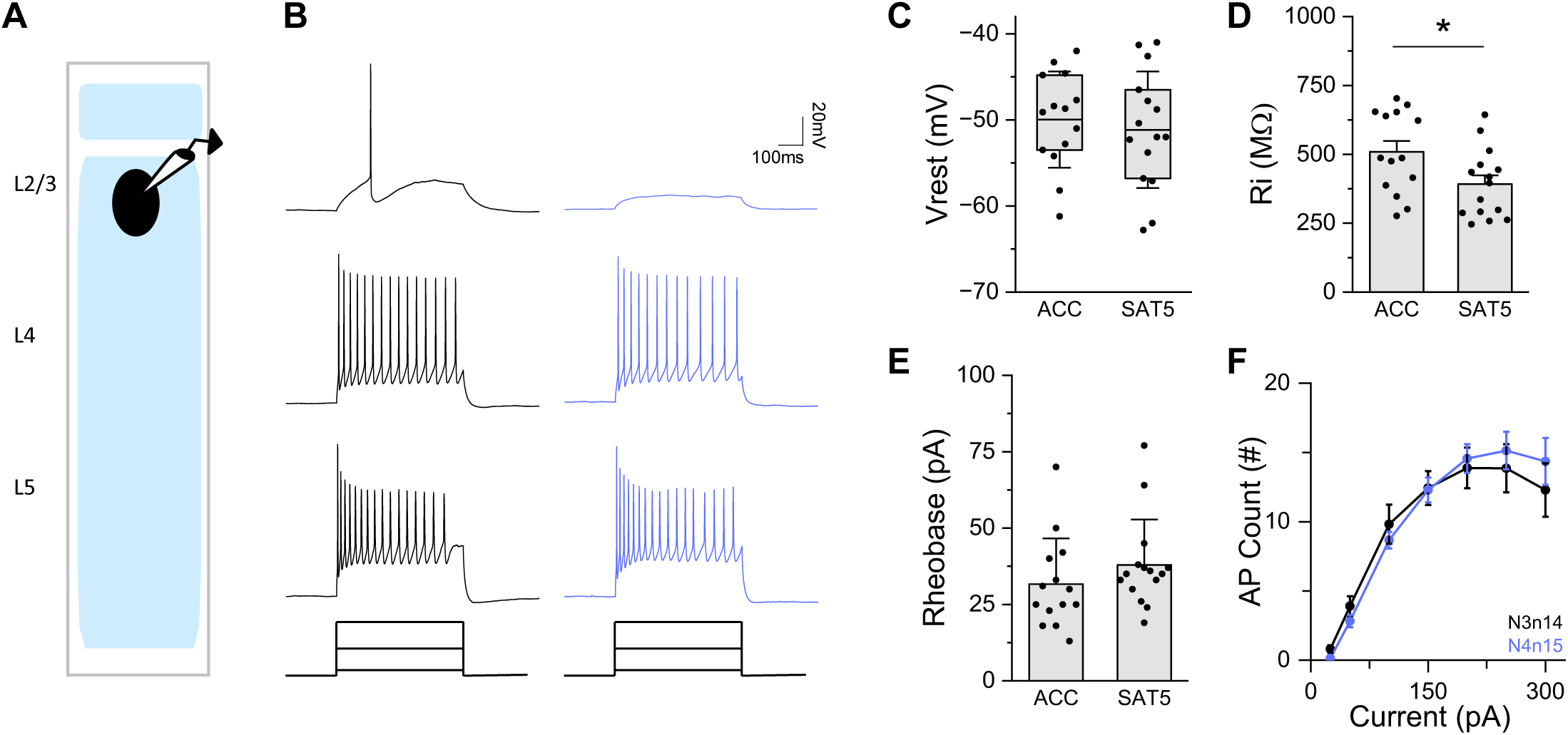
Intrinsic excitability of L2/3 SST neurons during SAT. (**A**) Schematic of the experiment setup. R2-expressing SST neurons were targeted. (**B**) Representative firing response following 500 ms 25 pA, 0 pA, and 300 pA current injection recorded from L2/3 low-threshold spiking SST neurons after 6 days acclimation (ACC6: black) and 5 days of SAT (SAT5: blue). (**C**) Resting membrane potential (Vrest) mparison. ACC6 (−50.0 ± 5.6 mV; N = 3 mice, n = 14 cells) vs. SAT5 (−51.1 ± 6.8 mV; N = 4 mice, n = cells). Box is 25th and 75th quartile, whiskers are SD, and midline is mean. Mann-Whitney U test (U =, n = 14 and 15, two-tailed). (**D**) Input resistance (Rin) comparison (mean + SEM). ACC6 (509.1 ± 39.8 Ω; N = 3 mice, n = 14 cells) vs. SAT5 (391.8 ± 32.0 MΩ; N = 4 mice, n = 15 cells), p=0.03. Mann-itney U test (U = 155, n = 14 and 15, two-tailed). (**E**) Rheobase comparison (mean + SD). ACC6 (31.6 5.0 pA, N = 3 mice, n = 14 cells) vs. SAT5 (37.9 ± 14.8 pA, N = 4 mice, n = 15 cells). Mann-Whitney est (U = 70.5, n = 15 and 14, two-tailed). (**F**) F-I curve of L2/3 SST neurons in ACC6 (black) and SAT5 ue) animals. Mean ± SEM. Two-way repeated measures ANOVA F(_1,27_) = 0.01, p = 0.91.

**Fig. S10.**
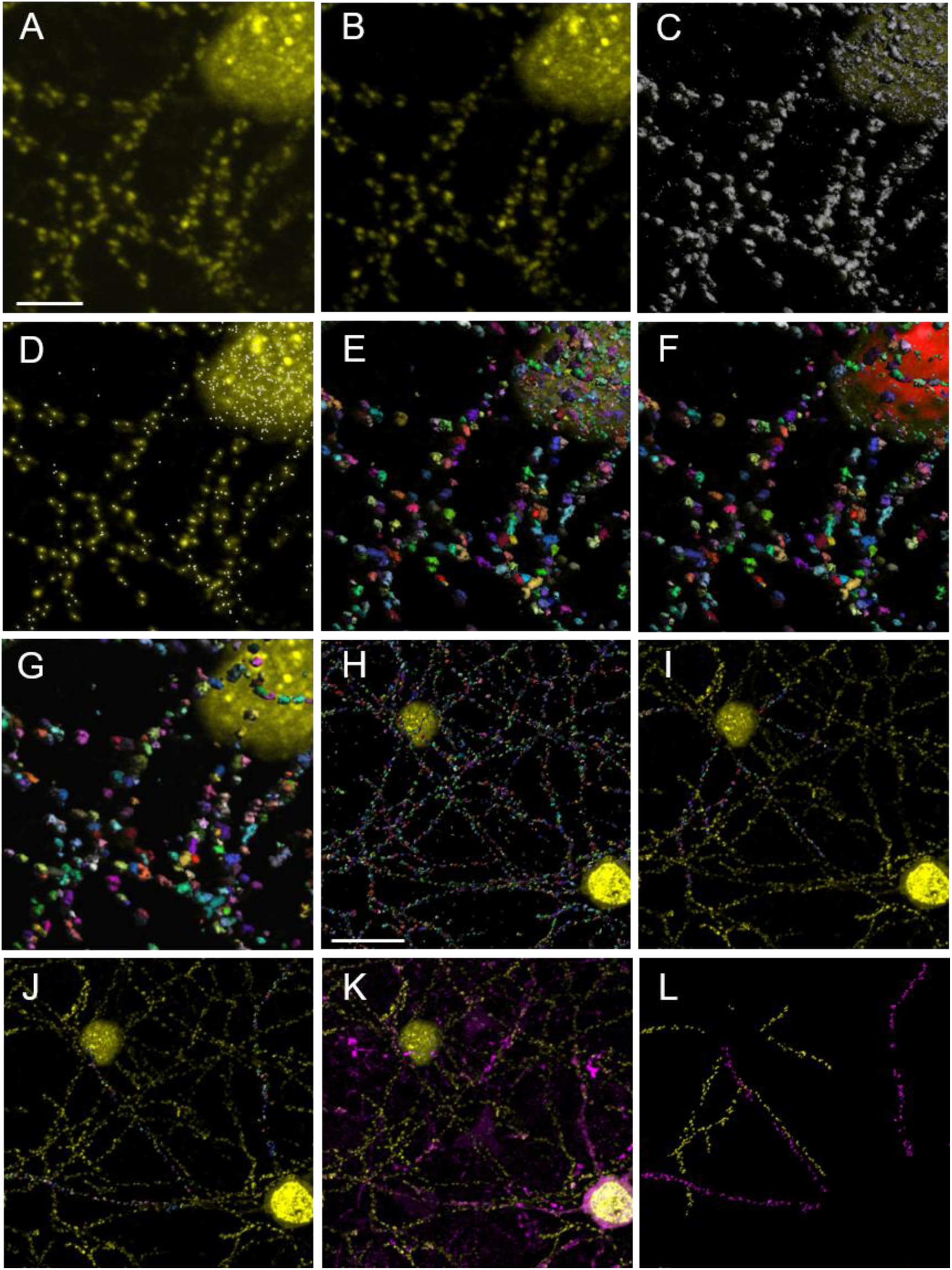
Analysis pipeline for PSD95-FingR puncta reconstruction and dendritic localization. (**A**) Region of interest (ROI) containing raw PSD95.FingR signal in confocal image stack (63x). Scale=5µm (**B**) Background subtraction prior surface reconstructions. (**C**) Semi-automated thresholding of 3D surface masks (gray) to optimize signal coverage while minimizing noise contained in background voxels. (**D**) A quality filter designating seed points (punctate signal) for splitting fused 3D surface masks. (**E**) 3D reconstructed fluorescence signal (multi-colored) after filtering structures to be ≥ 3 voxels to visualize all PSD95.FingR surface objects. (**F**) 3D surface mask of somatic PSD95.FingR signal. (**G**) 3D surface masks of PSD95.FingR signal within 0.15 um from reconstructed soma were removed. Remaining punctate PSD95.FingR signal (multi-colored) were included for whole field puncta analysis. (**H**) Zoomed out ROI from the same confocal image stack containing punctate PSD95.FingR signal for single cell puncta localization and subsequent comparisons between SST subtypes. Scale=15µm (**I**) Manually selected 3D surfaces (multi-colored) belonging to traceable dendrites emanating from soma in the top left corner. (**J**) Same as (I) but for soma in the bottom right corner. (**K**) Calb2-IR (purple) revealed for classifying puncta belonging to SST-Calb2 or SST-O. (**L**) Designated puncta belonging to SST-O (yellow) and SST-Calb2 (purple) neurons

**Fig. S11.**
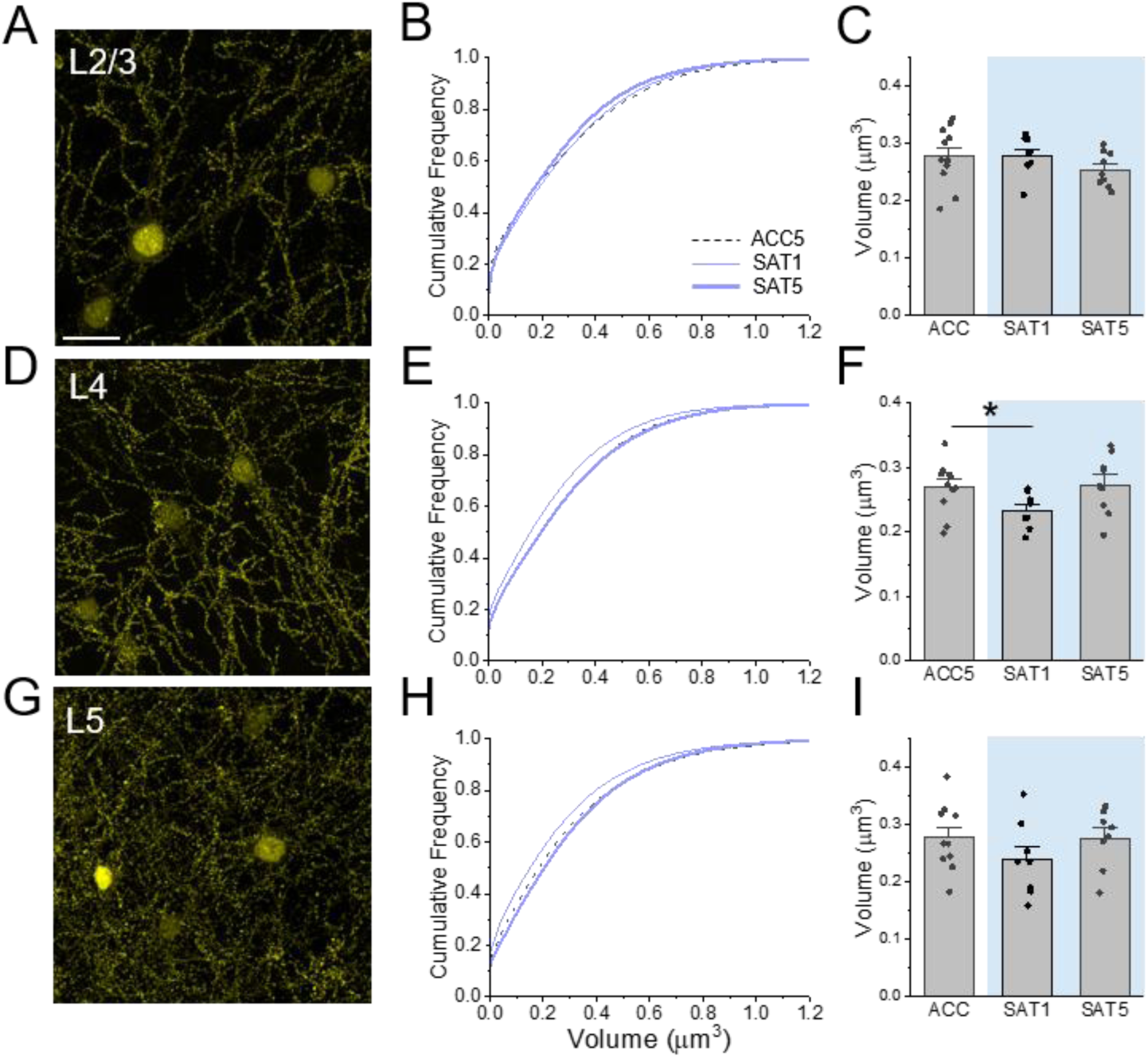
Field-of-view analysis of PSD95 puncta indicates layer-specific regulation of excitatory synapses onto SST neurons. (**A**) Representative field of view from PSD95.FingR labeled SST neurons in L2/3 of primary somatosensory cortex. Scale=20µm. (**B**) Cumulative distribution of reconstructed PSD95.FingR surface volume from whole field analysis in acclimated (black dashed line), one day (thin blue line), and five day trained (thick blue line) SST-Cre mice. (**C**) Mean puncta volume in individual animals across training conditions. (ACC N=11 mice, 60,500 puncta; SAT1 N=8 mice, 44,000 puncta; SAT5 N=9 mice, 49,500) puncta (**D-F**) Same as A-C but for L4 PSD95.FingR labeled SST neurons. (ACC N=11 mice, 79,420 puncta; SAT1 N=8 mice, 57,760 puncta; SAT5 N=9 mice, 64,980 puncta; ACC vs SAT1 p=.04, unpaired t-test) (**G-I**) Same as A-C but for L5 PSD95.FingR labeled SST neurons. (ACC N=11 mice, 98,900 puncta; SAT1 N=8 mice, 79,120 puncta; SAT5 N=9 mice, 89,010 puncta).

**Fig. S12.**
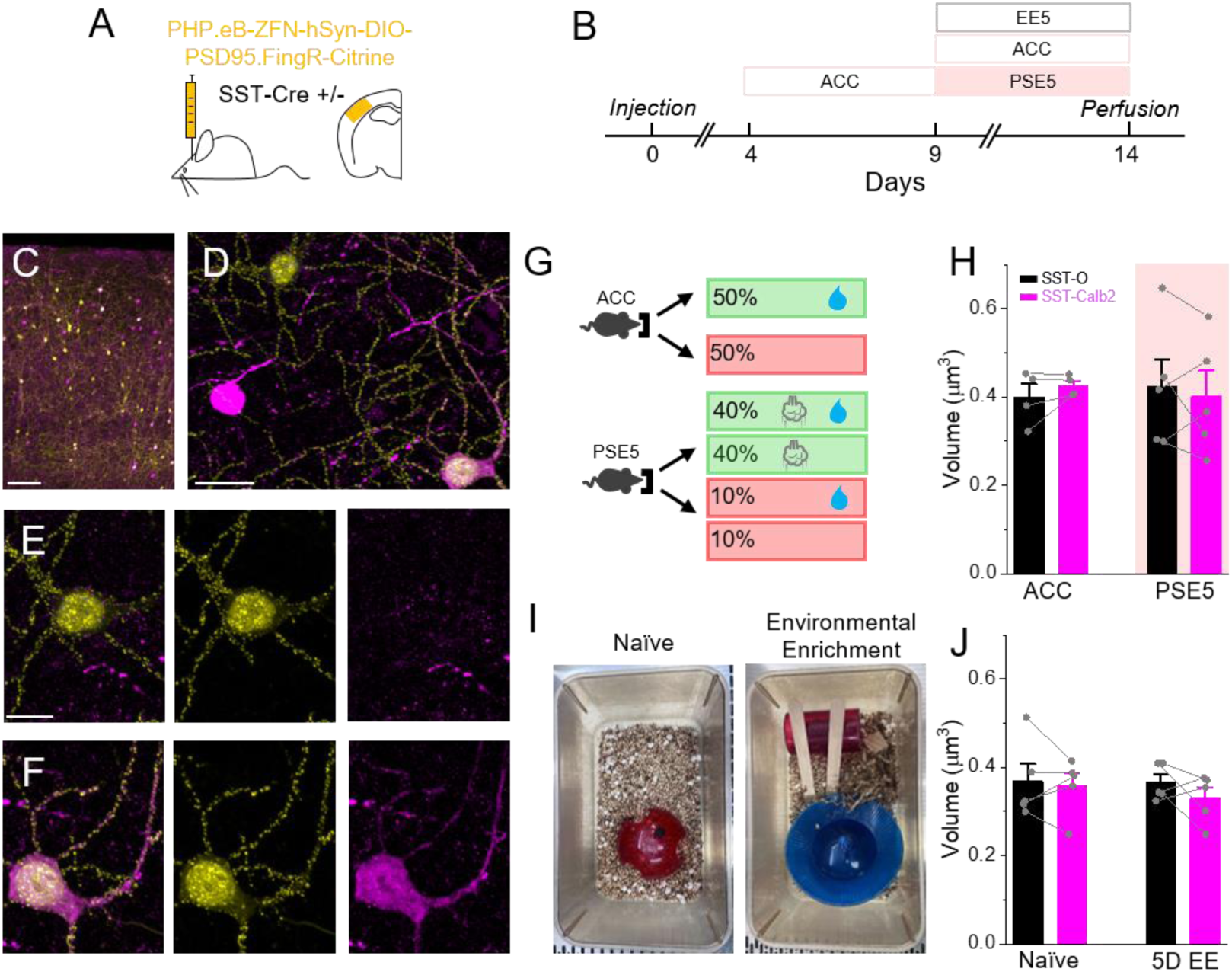
Neither pseudotraining nor sensory enrichment induce subtype-specific changes in L2/3 SST neurons. (**A**) Schematic outlining viral injection strategy. (**B**) Schematic outlining training timeline. (**C**) 10x image of PSD95.FingR labeled SST neurons merged with calretinin-IR. Scale = 100µm (**D**) 63x confocal image stack containing PSD95.FingR labeled SST neurons merged with calretinin-IR. Scale = 20µm (**E**) Zoomed image of SST-O neuron. PSD95.FingR-Citrine (left), calretinin-IR (middle), overlay, (right). Scale = 5µm (**F**) Same as (E) but for a SST-Calb2 neuron. (**G**) Schematic outlining stimulus and reward presentation probabilities after trial initiations in pseudo-training. (**H**) Within animal comparison between SST-O neurons (gray bars) and SST-Calb2 (pink bars) during acclimation or 5 days of pseudo-training (PSE5). (ACC N = 4mice, 800 puncta; PSE N = 5 mice, 1000 puncta). (**I**) Image showing naïve (left) and environmentally enriched cages (right). (**J**) Same as (H) but for naïve animals and animals experiencing five days of environmental enrichment (EE5). (Naive N = 5mice, 1000 puncta; EE5 N = 5 mice, 1000 puncta)

**Fig. S13.**
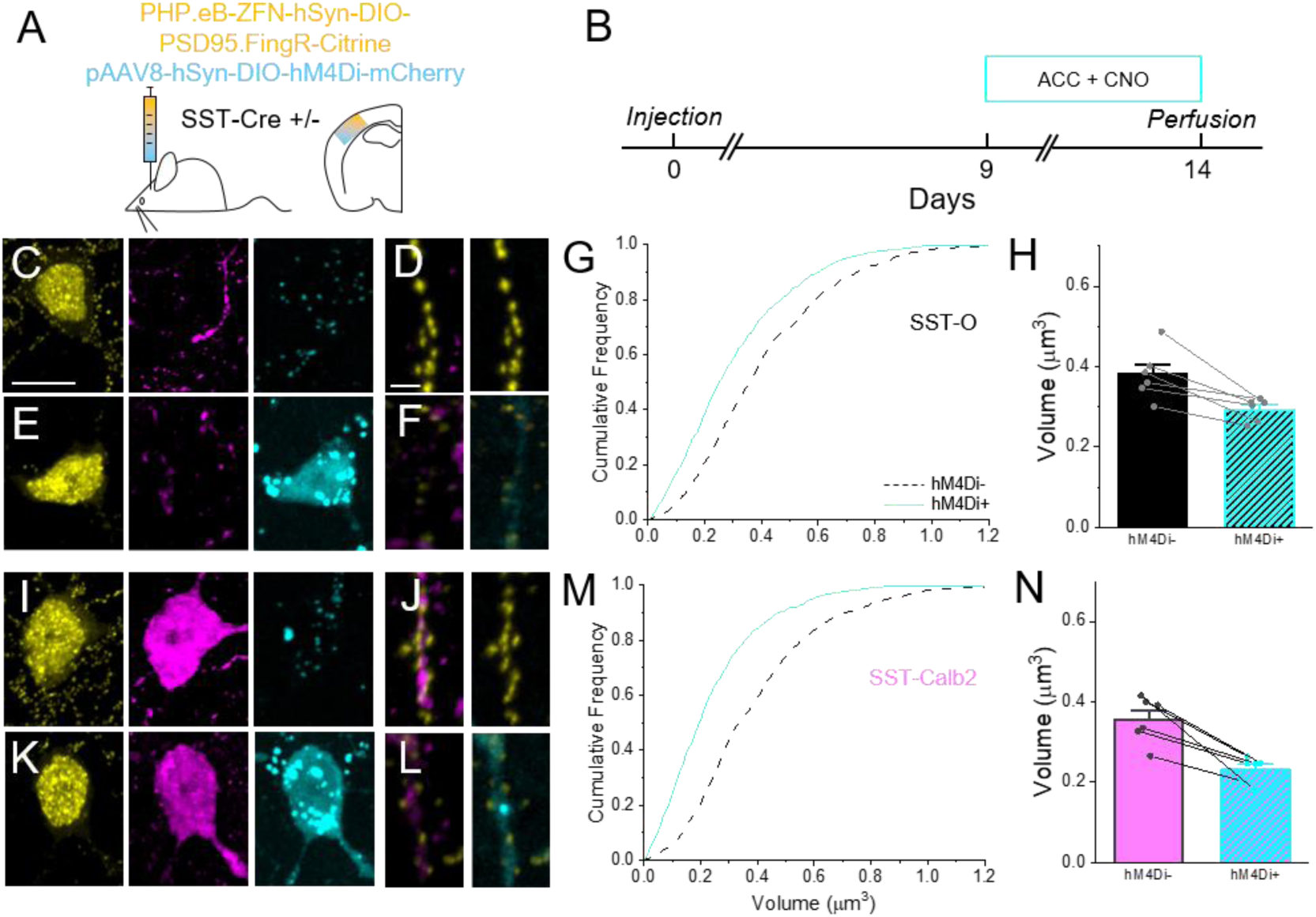
**Chemogenetic suppression of SST neuron activity reduces PSD95 puncta size on both SST-O and SST-Calb2 neurons**. (**A**) Schematic of Viral injection. (**B**) Schematic of experimental timeline. (**C**) 63x somatic ROI of volumetric confocal image stack containing from left to right, PSD95.FingR-Citrine (yellow), calretinin-IR (purple), and hM4Di-mCherry (pseudo-colored cyan) fluorescence channels. (**D**) Dendrite example from an SST-O neuron (left) overlay between PSD95.FingR (yellow) and calretinin-IR (purple). (right) overlay between PSD95.FingR and hM4Di-mCherry (cyan). (**E, F**) same as in (C, D) but for an SST-O neuron transduced with hM4Di-mCherry. (**G**) Cumulative distribution of PSD95 puncta volume in SST-O neurons with and without hM4Di expression (hM4Di-N=6 mice, 1200 puncta; hM4Di+ N=6 mice, 1200 puncta). (**H**) Within animal comparison of average PSD95 puncta size in SST-O cells with and without hM4Di expression. (**I, J**) same as in (C, D) but an SST-Calb2 neuron not transduced with hM4Di-mCherry. (**K, L**) same as in (C, D) but for an SST-Calb2 neuron transduced with hM4Di-mCherry. (**M, N**) Same as in (G,H) but for SST-Calb2 neurons (hM4Di-N=6 mice, 1200 puncta; hM4Di+ N=6 mice, 1200 puncta).

**Fig. S14:**
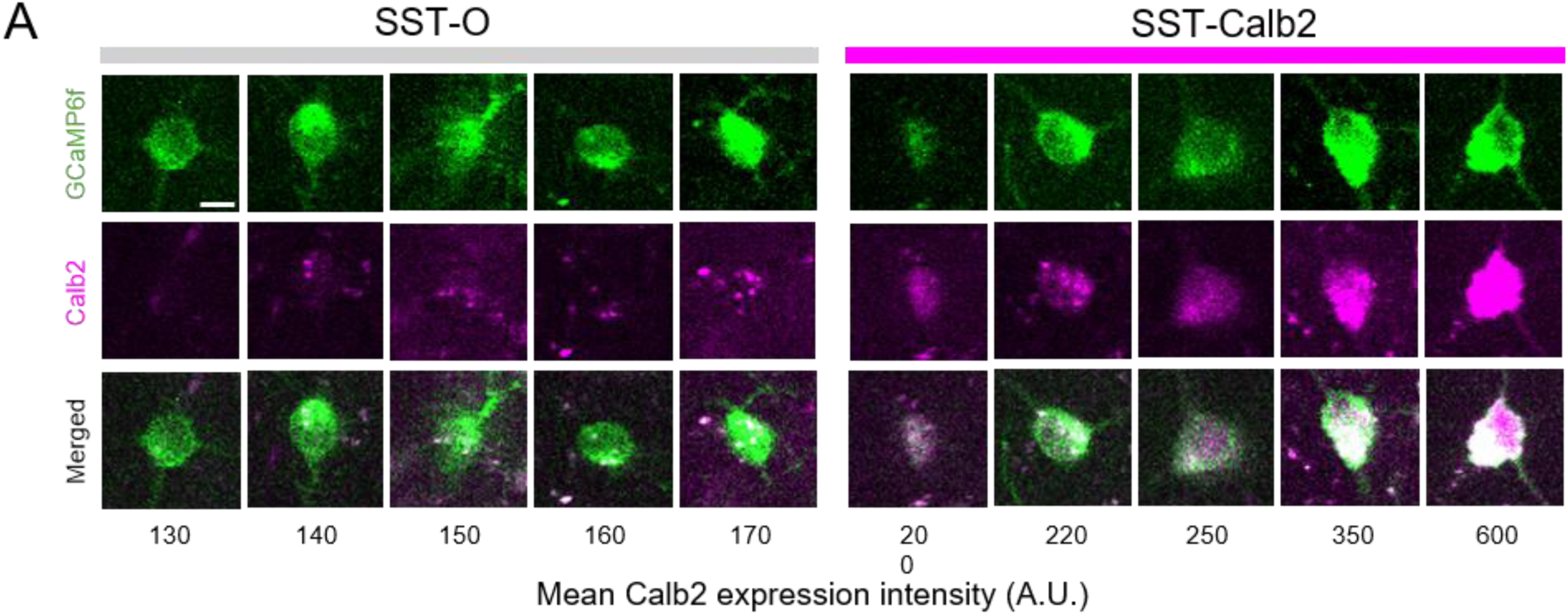
Range of Calb2-Cre reporter (mCherry) fluorescence intensity in L2/3 SST neurons. (**A**) Example neurons showing GCaMP6f expression (green) and Calb2 expression (magenta), with merged images (bottom) displaying both markers. The neurons are arranged from left to right in increasing order of Calb2 expression. Scale bar = 10μm.

**Fig. S15.**
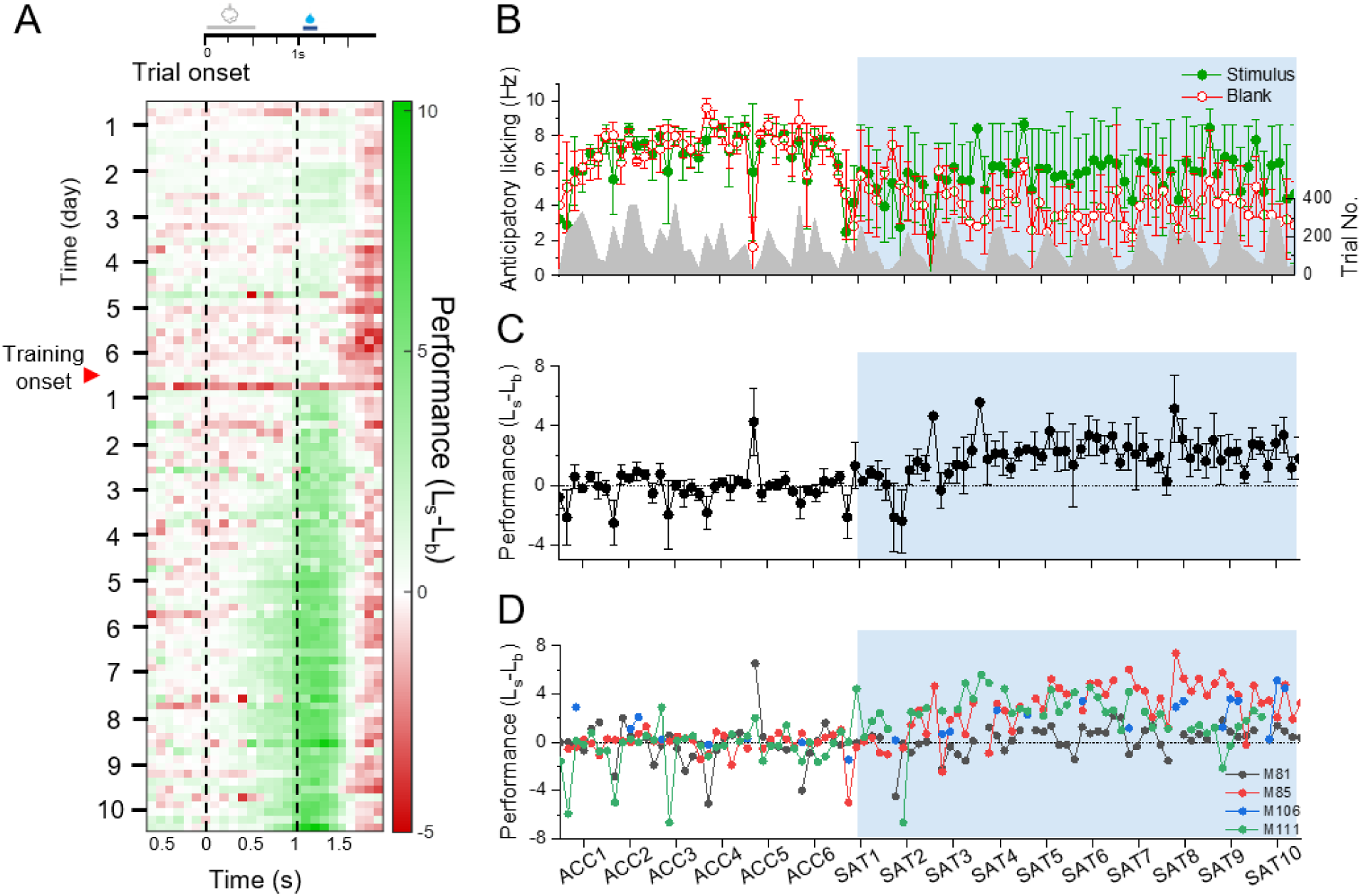
Changes in anticipatory licking in SST-Flp x Calb2-Cre transgenic mice during SAT. (**A**) Mean performance (Licking_stimulus_-Licking_blank_) averaged across 5 animals. (**B**) Mean anticipatory licking frequency averaged across 4 mice on stimulus (green) and blank (red) trials. (**C**) Mean performance averaged across 4 mice. (**D**) Performance of individual mice.

**Fig. S16.**
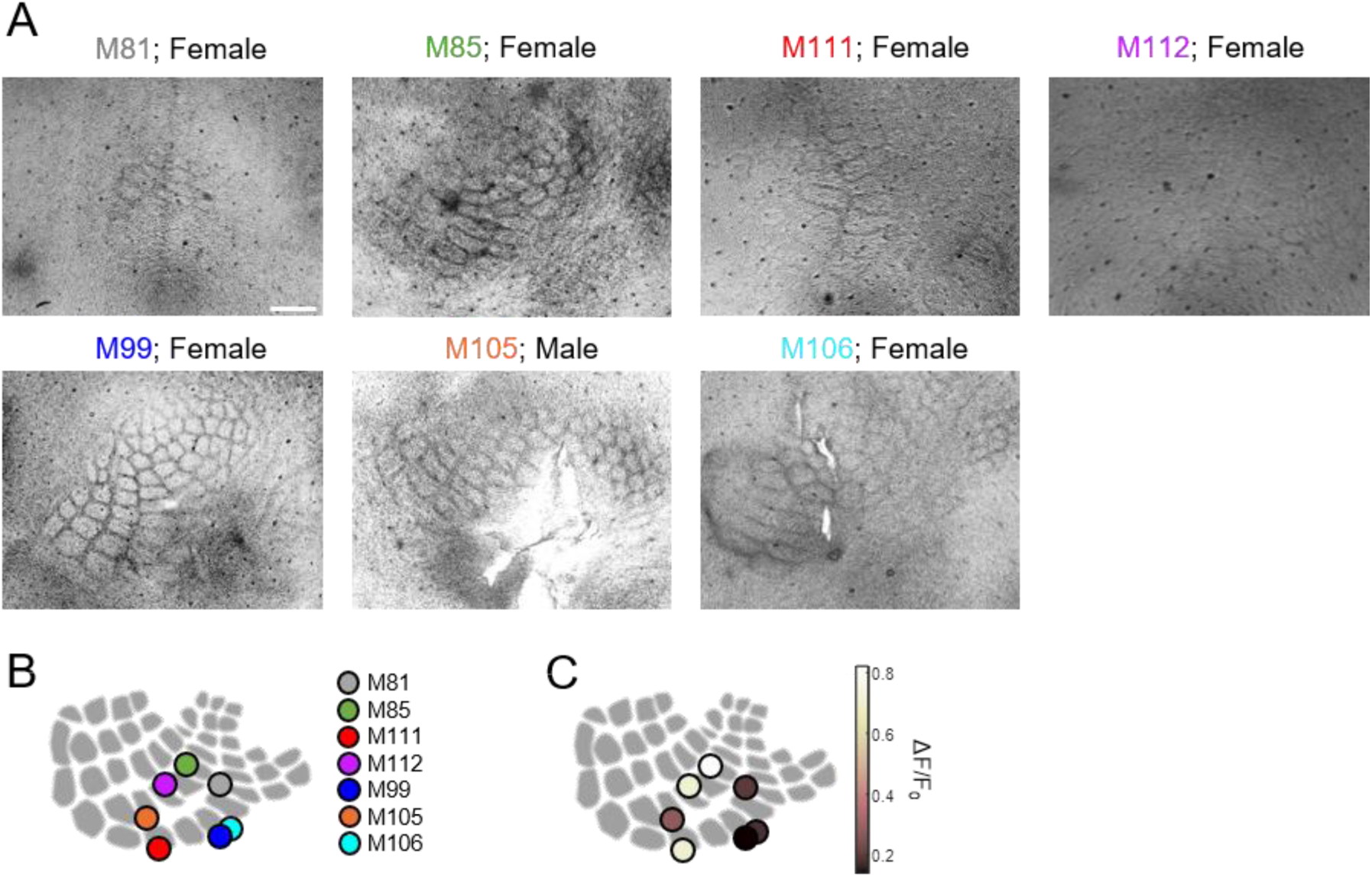
Post-hoc imaging location for SST-Flp x Calb2-Cre mice after SAT. (**A**) Post-hoc labeling of imaging site with a glass pipette filled with methyl blue dye. Scale=500 µm. (**B**) Schematic diagram of labeled imaging sites of all mice. Each dot represents a mouse. (**C**) Schematic diagram of labeled imaging sites of all mice color coded based on the mean peak response on ACC4-6.

**Fig. S17.**
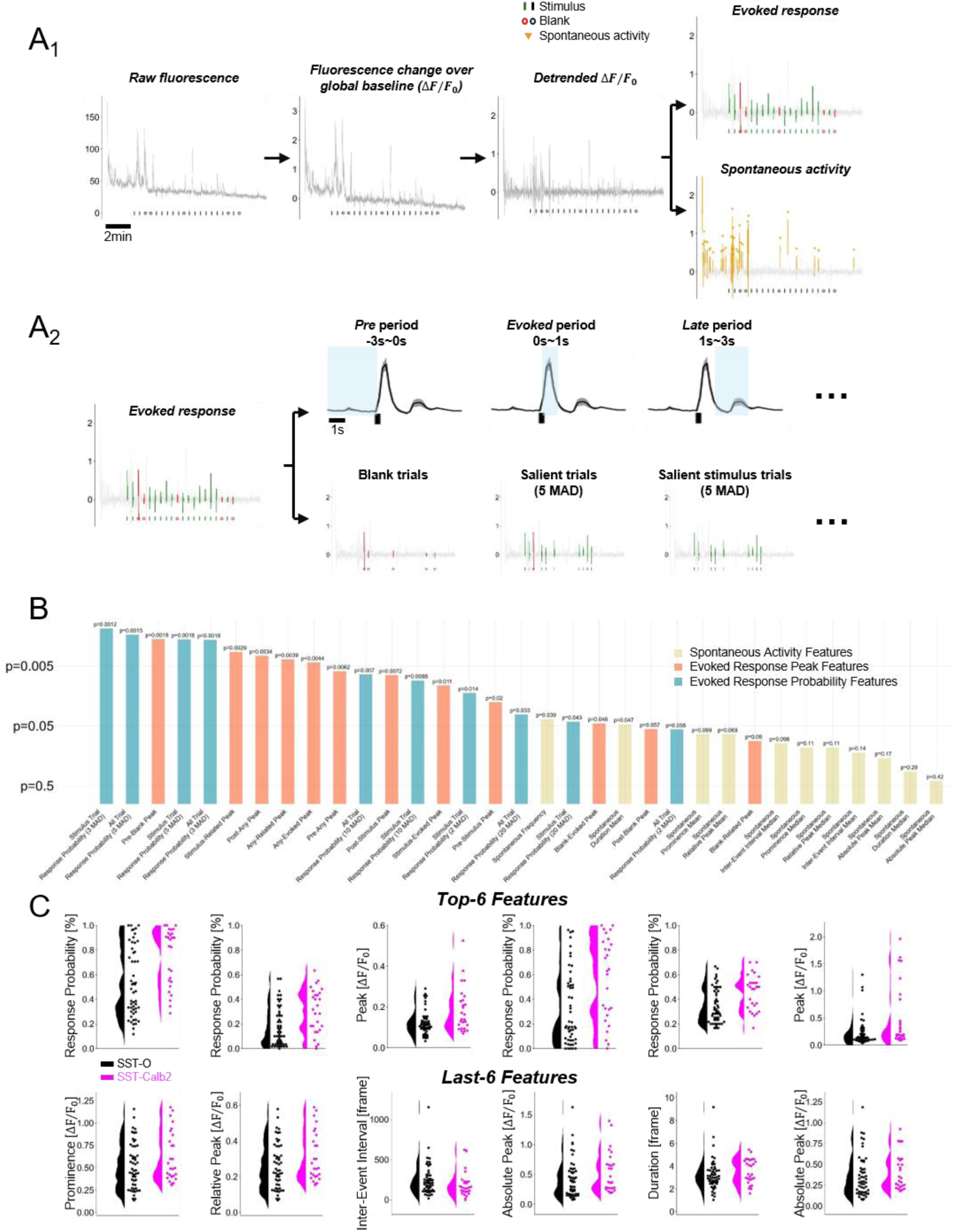
Extraction of response property features from whole-session recordings. (**A**) Feature extraction pipeline. (**A1**) Raw fluorescence signals were normalized to the global baseline and then detrended using a median filter (see Methods). After preprocessing, the detrended ΔF/F₀ signal was segmented into evoked responses and spontaneous activity. (**A2**) Example features extracted from evoked responses including evoked response peak features and evoked response probability features. (**B**) Ranking of all 33 features based on p-values from t-tests comparing SST-Calb2 and SST-O feature sets. (**C**) Example features showcasing the top 6 and bottom 6 ranked features from (B) averaged at ACC4-6.

**Fig. S18.**
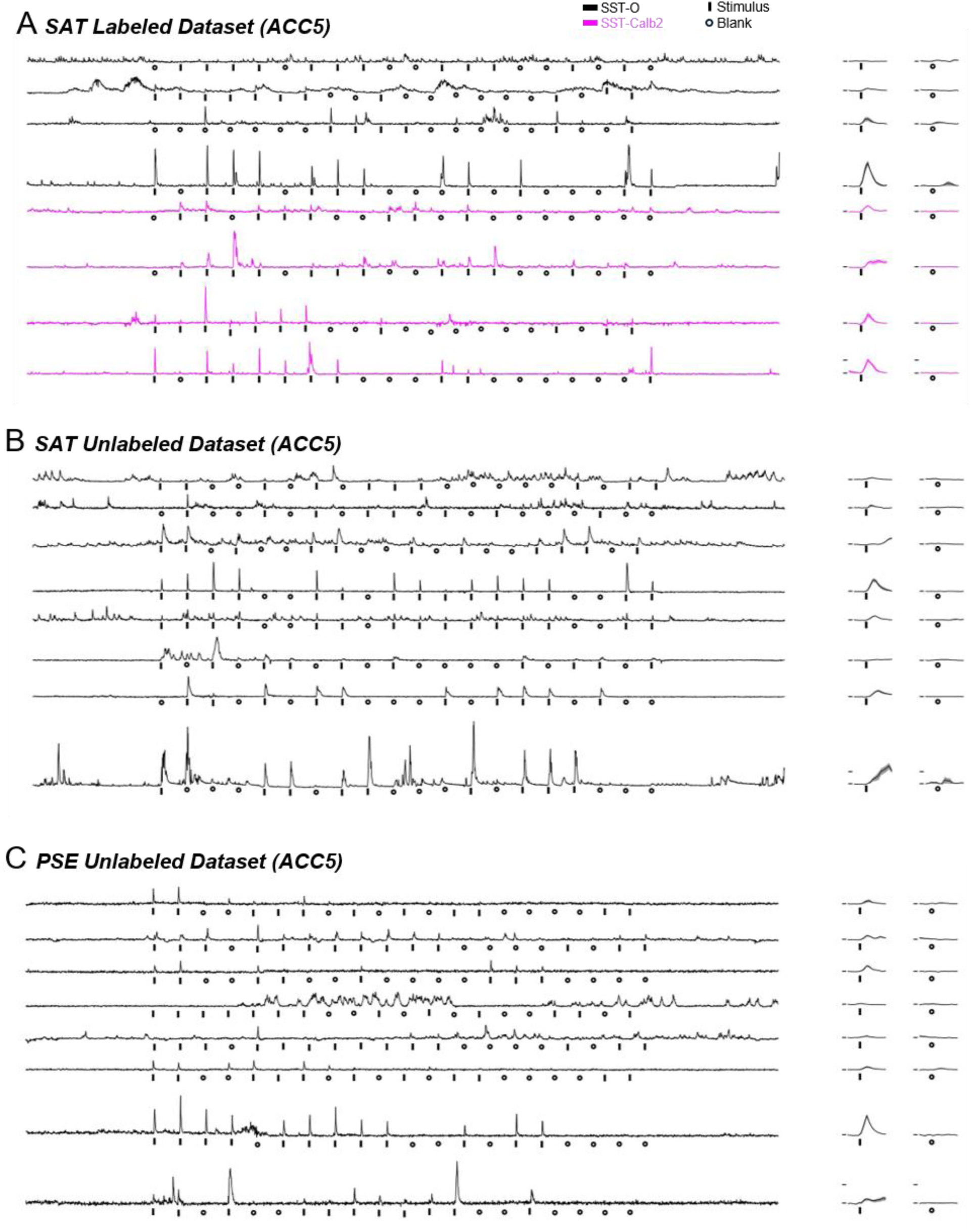
Example whole-session Ca^++^-imaging of L2/3 SST neurons. (**A**) Whole-session ACC5 example cells for SST-Flp x Calb2-iCre SAT dataset. (**B**) As in (A), but for SST-Cre x Ai148 SAT dataset. (**C**) As in (A), but for SST-Cre x Ai148 PSE dataset.

**Figure S19:**
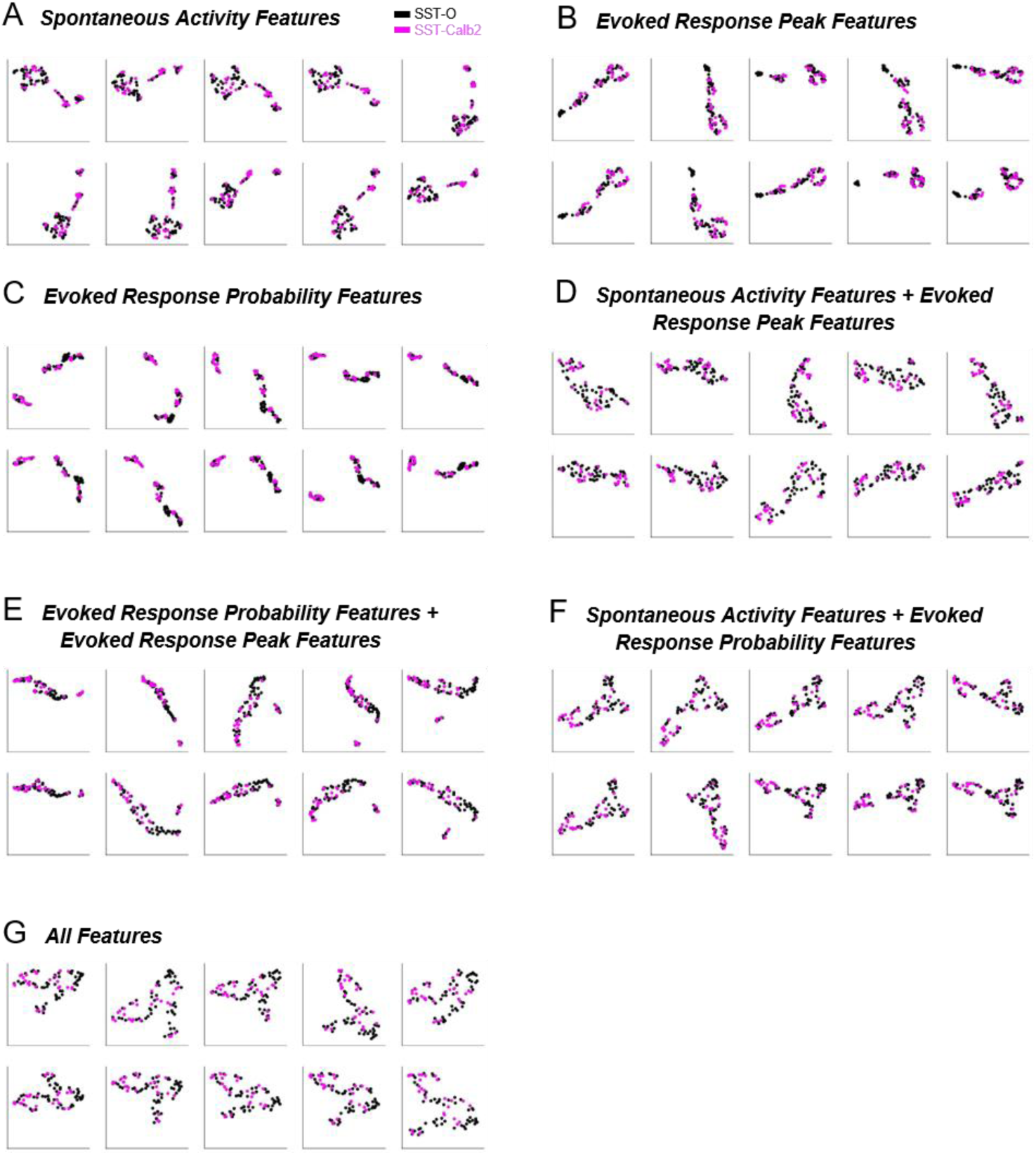
Robustness evaluation of different feature combinations for neuronal clustering. (**A-G**): UMAP visualizations of clustering results for SST-Flp x Calb2-iCre SAT dataset from ACC4-6, generated using 10 different random seeds to assess the robustness of the clustering method. Each panel represents the outcome of clustering based on distinct feature combinations

**Figure S20:**
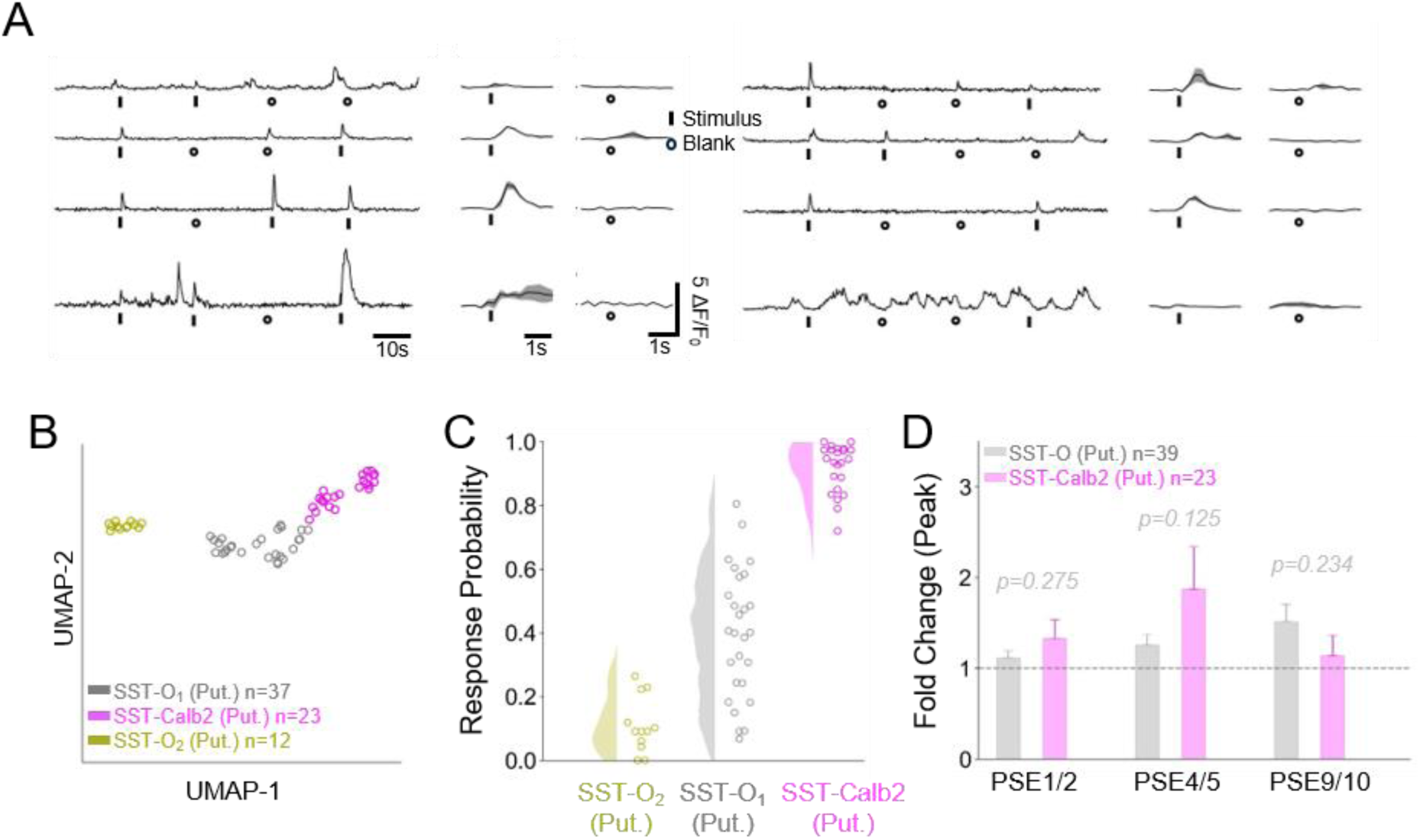
Classifier applied on PSE dataset predicts enhancement in both putative SST-Calb2 and SST-O cells. (**A-D**) as in Figure 4 (H-K), but for SST-Cre x Ai148 PSE dataset.

**Table S1.** Spontaneous Activity Features. List of spontaneous activity features from GCaMP6f fluorescence signals (See Methods)

| ***Feature Name*** | ***Description*** | ***Unit*** | ***Feature Name*** | ***Description*** | ***Unit*** |
| --- | --- | --- | --- | --- | --- |
| Absolute Peak Mean | mean of absolute detrended fluorescence of all spontaneous events | $\Delta F/F_0$ | Absolute Peak Median | median of absolute detrended fluorescence of all spontaneous events | $\Delta F/F_0$ |
| Prominence Mean | mean of vertical distance between the peak and its lowest contour line of all spontaneous events | $\Delta F/F_0$ | Prominence Median | median of vertical distance between the peak and its lowest contour line of all spontaneous events | $\Delta F/F_0$ |
| Inter-Event Interval Mean | mean of duration between adjacent events | *s* | Inter-Event Interval Median | median of duration between adjacent events | *s* |
| Duration Mean | mean of the lasting duration of all spontaneous events | *s* | Duration Median | median of the lasting duration of all spontaneous events | *s* |
| Relative Peak Mean | mean of the vertical distance between peak and its bases of all spontaneous events | $\Delta F/F_0$ | Relative Peak Median | median of the vertical distance between peak and its bases of all spontaneous events | $\Delta F/F_0$ |
| Frequency | number of spontaneous events per second | *Hz* |  |  |  |

**Table S2.** Evoked Response Peak Features. List of evoked response peak features from GCaMP6f fluorescence signals (See Methods)

| **Feature Name** | **Description** | **Unit** | **Feature Name** | **Description** | **Unit** | **Feature Name** | **Description** | **Unit** |
| --- | --- | --- | --- | --- | --- | --- | --- | --- |
| Pre-Stimulus Peak | mean of the peak detrended fluorescence during *pre* period (-3s~0s) of all stimulus trials | $\Delta F/F_0$ | Pre-Blank Peak | mean of the peak detrended fluorescence during *pre* period (-3s~0s) of all Blank trials | $\Delta F/F_0$ | Pre-Any Peak | mean of the peak detrended fluorescence during *pre* period (-3s~0s) of all trials | $\Delta F/F_0$ |
| Stimulus-Evoked Peak | mean of the peak detrended fluorescence during *evoked* period (0s~1s) of all stimulus trials | $\Delta F/F_0$ | Blank-Evoked Peak | mean of the peak detrended fluorescence during *evoked* period (0s~1s) of all Blank trials | $\Delta F/F_0$ | Any-Evoked Peak | mean of the peak detrended fluorescence during *evoked* period (0s~1s) of all trials | $\Delta F/F_0$ |
| Stimulus-Related Peak | mean of the peak detrended fluorescence during *late* period (1s~3s) of all stimulus trials | $\Delta F/F_0$ | Blank-Related Peak | mean of the peak detrended fluorescence during *late* period (1s~3s) of all Blank trials | $\Delta F/F_0$ | Any-Related Peak | mean of the peak detrended fluorescence during *late* period (1s~3s) of all trials | $\Delta F/F_0$ |
| Post-Stimulus Peak | mean of the peak detrended fluorescence during *post* period (0s~4s) of all stimulus trials | $\Delta F/F_0$ | Post-Blank Peak | mean of the peak detrended fluorescence during *post* period (0s~4s) of all Blank trials | $\Delta F/F_0$ | Post-Any Peak | mean of the peak detrended fluorescence during *post* period (0s~4s) of all trials | $\Delta F/F_0$ |

**Table S3.** Evoked Response Probability Features. List of evoked response probability features from GCaMP6f fluorescence signals (See Methods).

| *Feature Name* | *Description* | *Unit* | *Feature Name* | *Description* | *Unit* |
| --- | --- | --- | --- | --- | --- |
| Stimulus Trial Response Probability with threshold 2 | fraction of stimulus trial whose *evoked* period (0s~1s) peak larger or equal to 2 times the median absolute deviation (MAD) | % | All Trial Response Probability with threshold 2 | fraction of any trial whose *evoked* period (0s~1s) peak larger or equal to 2 times the median absolute deviation (MAD) | % |
| Stimulus Trial Response Probability with threshold 3 | fraction of stimulus trial whose *evoked* period (0s~1s) peak larger or equal to 3 times the median absolute deviation (MAD) | % | All Trial Response Probability with threshold 3 | fraction of any trial whose *evoked* period (0s~1s) peak larger or equal to 3 times the median absolute deviation (MAD) | % |
| Stimulus Trial Response Probability with threshold 5 | fraction of stimulus trial whose *evoked* period (0s~1s) peak larger or equal to 5 times the median absolute deviation (MAD) | % | All Trial Response Probability with threshold 5 | fraction of any trial whose *evoked* period (0s~1s) peak larger or equal to 5 times the median absolute deviation (MAD) | % |
| Stimulus Trial Response Probability with threshold 10 | fraction of stimulus trial whose *evoked* period (0s~1s) peak larger or equal to 10 times the median absolute deviation (MAD) | % | All Trial Response Probability with threshold 10 | fraction of any trial whose *evoked* period (0s~1s) peak larger or equal to 10 times the median absolute deviation (MAD) | % |
| Stimulus Trial Response Probability with threshold 20 | fraction of stimulus trial whose *evoked* period (0s~1s) peak larger or equal to 20 times the median absolute deviation (MAD) | % | All Trial Response Probability with threshold 20 | fraction of any trial whose *evoked* period (0s~1s) peak larger or equal to 20 times the median absolute deviation (MAD) | % |

## References and Notes

1. Z. Yao, C. T. J. van Velthoven, M. Kunst, M. Zhang, D. McMillen, C. Lee, W. Jung, J. Goldy, A. Abdelhak, M. Aitken, K. Baker, P. Baker, E. Barkan, D. Bertagnolli, A. Bhandiwad, C. Bielstein, P. Bishwakarma, J. Campos, D. Carey, T. Casper, A. B. Chakka, R. Chakrabarty, S. Chavan, M. Chen, M. Clark, J. Close, K. Crichton, S. Daniel, P. DiValentin, T. Dolbeare, L. Ellingwood, E. Fiabane, T. Fliss, J. Gee, J. Gerstenberger, A. Glandon, J. Gloe, J. Gould, J. Gray, N. Guilford, J. Guzman, D. Hirschstein, W. Ho, M. Hooper, M. Huang, M. Hupp, K. Jin, M. Kroll, K. Lathia, A. Leon, S. Li, B. Long, Z. Madigan, J. Malloy, J. Malone, Z. Maltzer, N. Martin, R. McCue, R. McGinty, N. Mei, J. Melchor, E. Meyerdierks, T. Mollenkopf, S. Moonsman, T. N. Nguyen, S. Otto, T. Pham, C. Rimorin, A. Ruiz, R. Sanchez, L. Sawyer, N. Shapovalova, N. Shepard, C. Slaughterbeck, J. Sulc, M. Tieu, A. Torkelson, H. Tung, N. Valera Cuevas, S. Vance, K. Wadhwani, K. Ward, B. Levi, C. Farrell, R. Young, B. Staats, M.-Q. M. Wang, C. L. Thompson, S. Mufti, C. M. Pagan, L. Kruse, N. Dee, S. M. Sunkin, L. Esposito, M. J. Hawrylycz, J. Waters, L. Ng, K. Smith, B. Tasic, X. Zhuang, H. Zeng, A high-resolution transcriptomic and spatial atlas of cell types in the whole mouse brain. Nature 624, 317– 332 (2023).

2. A. Agmon, A. L. Barth, A brief history of somatostatin interneuron taxonomy or: how many somatostatin subtypes are there, really? *Front*. Neural Circuits 18 (2024).

3. H. Adesnik, W. Bruns, H. Taniguchi, Z. J. Huang, M. Scanziani, A neural circuit for spatial summation in visual cortex. Nature 490, 226–231 (2012).

4. J. P. Hamm, R. Yuste, Somatostatin Interneurons Control a Key Component of Mismatch Negativity in Mouse Visual Cortex. Cell Reports 16, 597–604 (2016).

5. J. Poort, K. A. Wilmes, A. Blot, A. Chadwick, M. Sahani, C. Clopath, T. D. Mrsic-Flogel, S. B. Hofer, A. G. Khan, Learning and attention increase visual response selectivity through distinct mechanisms. Neuron 110, 686–697.e6 (2022).

6. H. Makino, T. Komiyama, Learning enhances the relative impact of top-down processing in the visual cortex. Nat Neurosci 18, 1116–1122 (2015).

7. J. Yang, P. Serrano, X. Yin, X. Sun, Y. Lin, S. X. Chen, Functionally distinct NPAS4-expressing somatostatin interneuron ensembles critical for motor skill learning. Neuron 110, 3339–3355.e8 (2022).

8. S. Furutachi, A. D. Franklin, A. M. Aldea, T. D. Mrsic-Flogel, S. B. Hofer, Cooperative thalamocortical circuit mechanism for sensory prediction errors. Nature, 1–9 (2024).

9. A. Adler, R. Zhao, M. E. Shin, R. Yasuda, W.-B. Gan, Somatostatin-Expressing Interneurons Enable and Maintain Learning-Dependent Sequential Activation of Pyramidal Neurons. Neuron 102, 202–216.e7 (2019).

10. G. Dobrzanski, A. Lukomska, R. Zakrzewska, A. Posluszny, D. Kanigowski, J. Urban-Ciecko, M. Liguz-Lecznar, M. Kossut, Learning-induced plasticity in the barrel cortex is disrupted by inhibition of layer 4 somatostatin-containing interneurons. Biochim Biophys Acta Mol Cell Res 1869, 119146 (2022).

11. J. Yu, H. Hu, A. Agmon, K. Svoboda, Recruitment of GABAergic Interneurons in the Barrel Cortex during Active Tactile Behavior. Neuron 104, 412–427.e4 (2019).

12. M. de Brito Van Velze, D. Dhanasobhon, M. Martinez, A. Morabito, E. Berthaux, C. M. Pinho, Y. Zerlaut, N. Rebola, Feedforward and disinhibitory circuits differentially control activity of cortical somatostatin interneurons during behavioral state transitions. Cell Rep 43, 114197 (2024).

13. H. K. Kato, S. N. Gillet, J. S. Isaacson, Flexible Sensory Representations in Auditory Cortex Driven by Behavioral Relevance. Neuron 88, 1027–1039 (2015).

14. N. J. Audette, S. M. Bernhard, A. Ray, L. T. Stewart, A. L. Barth, Rapid Plasticity of Higher-Order Thalamocortical Inputs during Sensory Learning. Neuron 103, 277–291.e4 (2019).

15. J. Guy, M. Möck, J. F. Staiger, Direction selectivity of inhibitory interneurons in mouse barrel cortex differs between interneuron subtypes. Cell Rep 42, 111936 (2023).

16. C. F. Khoury, N. G. Fala, C. A. Runyan, Arousal and Locomotion Differently Modulate Activity of Somatostatin Neurons across Cortex. eNeuro 10 (2023).

17. C. Condylis, E. Lowet, J. Ni, K. Bistrong, T. Ouellette, N. Josephs, J. L. Chen, Context-Dependent Sensory Processing across Primary and Secondary Somatosensory Cortex. Neuron 106, 515–525.e5 (2020).

18. W. Muñoz, R. Tremblay, D. Levenstein, B. Rudy, Layer-specific modulation of neocortical dendritic inhibition during active wakefulness. Science 355, 954–959 (2017).

19. A. Ray, J. A. Christian, M. B. Mosso, E. Park, W. Wegner, K. I. Willig, A. L. Barth, Quantitative Fluorescence Analysis Reveals Dendrite-Specific Thalamocortical Plasticity in L5 Pyramidal Neurons during Learning. J Neurosci 43, 584–600 (2023).

20. J. Urban-Ciecko, A. L. Barth, Somatostatin-expressing neurons in cortical networks. Nat. Rev. Neurosci. 17, 401–409 (2016).

21. A. Nowacka, M. Borczyk, A. Salamian, T. Wójtowicz, J. Włodarczyk, K. Radwanska, PSD-95 Serine 73 phosphorylation is not required for induction of NMDA-LTD. Sci Rep 10, 2054 (2020).

22. G. G. Gross, J. A. Junge, R. J. Mora, H.-B. Kwon, C. A. Olson, T. T. Takahashi, E. R. Liman, G. C. R. Ellis-Davies, A. W. McGee, B. L. Sabatini, R. W. Roberts, D. B. Arnold, Recombinant probes for visualizing endogenous synaptic proteins in living neurons. Neuron 78, 971–985 (2013).

23. N. W. Gouwens, S. A. Sorensen, F. Baftizadeh, A. Budzillo, B. R. Lee, T. Jarsky, L. Alfiler, K. Baker, E. Barkan, K. Berry, D. Bertagnolli, K. Bickley, J. Bomben, T. Braun, K. Brouner, T. Casper, K. Crichton, T. L. Daigle, R. Dalley, R. A. de Frates, N. Dee, T. Desta, S. D. Lee, N. Dotson, T. Egdorf, L. Ellingwood, R. Enstrom, L. Esposito, C. Farrell, D. Feng, O. Fong, R. Gala, C. Gamlin, A. Gary, A. Glandon, J. Goldy, M. Gorham, L. Graybuck, H. Gu, K. Hadley, M. J. Hawrylycz, A. M. Henry, D. Hill, M. Hupp, S. Kebede, T. K. Kim, L. Kim, M. Kroll, C. Lee, K. E. Link, M. Mallory, R. Mann, M. Maxwell, M. McGraw, D. McMillen, A. Mukora, L. Ng, L. Ng, K. Ngo, P. R. Nicovich, A. Oldre, D. Park, H. Peng, O. Penn, T. Pham, A. Pom, Z. Popović, L. Potekhina, R. Rajanbabu, S. Ransford, D. Reid, C. Rimorin, M. Robertson, K. Ronellenfitch, A. Ruiz, D. Sandman, K. Smith, J. Sulc, S. M. Sunkin, A. Szafer, M. Tieu, A. Torkelson, J. Trinh, H. Tung, W. Wakeman, K. Ward, G. Williams, Z. Zhou, J. T. Ting, A. Arkhipov, U. Sümbül, E. S. Lein, C. Koch, Z. Yao, B. Tasic, J. Berg, G. J. Murphy, H. Zeng, Integrated Morphoelectric and Transcriptomic Classification of Cortical GABAergic Cells. Cell 183, 935–953.e19 (2020).

24. S. J. Wu, E. Sevier, D. Dwivedi, G.-A. Saldi, A. Hairston, S. Yu, L. Abbott, D. H. Choi, M. Sherer, Y. Qiu, A. Shinde, M. Lenahan, D. Rizzo, Q. Xu, I. Barrera, V. Kumar, G. Marrero, A. Prönneke, S. Huang, K. Kullander, D. A. Stafford, E. Macosko, F. Chen, B. Rudy, G. Fishell, Cortical somatostatin interneuron subtypes form cell-type-specific circuits. Neuron 111, 2675–2692.e9 (2023).

25. R. E. Hostetler, H. Hu, A. Agmon, Genetically Defined Subtypes of Somatostatin-Containing Cortical Interneurons. eNeuro 10, ENEURO.0204-23.2023 (2023).

26. R. G. Northcutt, J. H. Kaas, The emergence and evolution of mammalian neocortex. Trends Neurosci 18, 373–379 (1995).

27. C. M. Niell, M. Scanziani, How Cortical Circuits Implement Cortical Computations: Mouse Visual Cortex as a Model. Annu Rev Neurosci 44, 517–546 (2021).

28. X. Xu, E. M. Callaway, Laminar specificity of functional input to distinct types of inhibitory cortical neurons. J Neurosci 29, 70–85 (2009).

29. A. Attinger, B. Wang, G. B. Keller, Visuomotor Coupling Shapes the Functional Development of Mouse Visual Cortex. Cell 169, 1291–1302.e14 (2017).

30. C. K. Pfeffer, M. Xue, M. He, Z. J. Huang, M. Scanziani, Inhibition of inhibition in visual cortex: the logic of connections between molecularly distinct interneurons. Nat. Neurosci. 16, 1068–1076 (2013).

31. J. U. Henschke, E. Dylda, D. Katsanevaki, N. Dupuy, S. P. Currie, T. Amvrosiadis, J. M. P. Pakan, N. L. Rochefort, Reward Association Enhances Stimulus-Specific Representations in Primary Visual Cortex. Curr Biol 30, 1866–1880.e5 (2020).

32. A. Schneider, M. Azabou, L. McDougall-Vigier, D. F. Parks, S. Ensley, K. Bhaskaran-Nair, T. Nowakowski, E. L. Dyer, K. B. Hengen, Transcriptomic cell type structures in vivo neuronal activity across multiple timescales. Cell Rep 42, 112318 (2023).

33. C. Trainito, C. von Nicolai, E. K. Miller, M. Siegel, Extracellular Spike Waveform Dissociates Four Functionally Distinct Cell Classes in Primate Cortex. Curr Biol 29, 2973–2982.e5 (2019).

34. S. M. Bernhard, J. Lee, M. Zhu, A. Hsu, A. Erskine, S. A. Hires, A. L. Barth, An automated homecage system for multiwhisker detection and discrimination learning in mice. PLoS One 15, e0232916 (2020).

35. M. Zhu, S. J. Kuhlman, A. L. Barth, Transient enhancement of stimulus-evoked activity in neocortex during sensory learning. Learn Mem 31, a053870 (2024).

36. M. Pachitariu, C. Stringer, M. Dipoppa, S. Schröder, L. F. Rossi, H. Dalgleish, M. Carandini, K. D. Harris, Suite2p: beyond 10,000 neurons with standard two-photon microscopy. bioRxiv [Preprint] (2017). 10.1101/061507.

37. T.-W. Chen, T. J. Wardill, Y. Sun, S. R. Pulver, S. L. Renninger, A. Baohan, E. R. Schreiter, R. A. Kerr, M. B. Orger, V. Jayaraman, L. L. Looger, K. Svoboda, D. S. Kim, Ultrasensitive fluorescent proteins for imaging neuronal activity. Nature 499, 295–300 (2013).

